# Detection of Spacer Precursors Formed *In Vivo* During Primed CRISPR Adaptation

**DOI:** 10.1101/594259

**Authors:** Anna A. Shiriaeva, Ekaterina Savitskaya, Kirill A. Datsenko, Irina O. Vvedenskaya, Iana Fedorova, Natalia Morozova, Anastasia Metlitskaya, Anton Sabantsev, Bryce E. Nickels, Konstantin Severinov, Ekaterina Semenova

**Affiliations:** Center for Data-Intensive Biomedicine and Biotechnology, Skolkovo Institute of Science and Technology, Moscow, 143028, Russia; Peter the Great St. Petersburg Polytechnic University, St. Petersburg, Russia, 195251; Department of Molecular Biology and Biochemistry, Waksman Institute, Rutgers University, Piscataway, NJ 08854, USA; Institute of Molecular Genetics, Russian Academy of Sciences, Moscow, 123182; Department of Genetics and Waksman Institute, Rutgers University, Piscataway, NJ 08854, USA

## Abstract

Type I CRISPR (Clustered Regularly Interspaced Short Palindromic Repeats)-Cas (CRISPR associated) loci provide prokaryotes with a nucleic-acid-based adaptive immunity against foreign DNA^1^. Immunity involves “adaptation,” the integration of ~30-bp DNA fragments into the CRISPR array as “spacer” sequences, and “interference,” the targeted degradation of DNA containing a “protospacer” sequence mediated by a complex containing a spacer-derived CRISPR RNA (crRNA)^1–4^. Specificity for targeting interference to protospacers, but not spacers, occurs through recognition of a 3-bp protospacer adjacent motif (PAM)^5^ by the crRNA-containing complex^6^. Interference-driven DNA degradation of protospacer-containing DNA can be coupled with “primed adaptation,” ill which spacers are acquired from DNA surrounding the targeted protospacer in a bidirectional, orientation-dependent manner^2,3,7^. Here we construct a robust *in vivo* model for primed adaptation consisting of an *Escherichia coli* type I-E CRISPR-Cas “self-targeting” locus encoding a crRNA that targets a chromosomal protospacer. We develop a strand-specific, high-throughput-sequencing method for analysis of DNA fragments, “FragSeq,” and use this method to detect short fragments derived from DNA surrounding the targeted protospacer. The detected fragments have sequences matching spacers acquired during primed adaptation, contain ~3- to 4-nt overhangs derived from excision of genomic DNA within a PAM, are generated in a bidirectional, orientation-dependent manner relative to the targeted protospacer, require the functional integrity of machinery for interference and adaptation to accumulate, and function as spacer precursors when exogenously introduced into cells by transformation. DNA fragments with a similar structure accumulate in cells undergoing primed adaptation in a type I-F CRISPR-Cas self-targeting system. We propose the DNA fragments detected in this work are products of universal steps of spacer precursor processing in type I CRISPR-Cas systems.

---

CRISPR interference in the *E. coli* type I-E system is performed by the Cascade complex, composed of a crRNA and several Cas proteins^1,8,9^. Initial binding of Cascade to a protospacer results in formation of an R-loop containing an RNA-DNA heteroduplex formed between the crRNA and “target” strand, and extrusion of single-stranded DNA derived from the “nontarget” strand^6,8,10–14^. Cas3, a single-stranded nuclease and 3’-5’ helicase, is recruited to the Cascade-protospacer complex and cleaves the nontarget strand to initiate unwinding and degradation of the targeted DNA^11,13,15^. *In vitro*, Cas3 can translocate on DNA as a component of a larger complex that includes Cascade and the key proteins of CRISPR adaptation, Cas1 and Cas2^16^.

CRISPR adaptation in the *E. coli* I-E system is mediated by a Cas1-Cas2 complex that can facilitate spacer acquisition in the absence of interference, a process termed “naive adaptation”^14,17–19^. In primed CRISPR adaptation, interference-driven DNA degradation initiated at a “priming protospacer (PPS),” is coupled with acquisition of spacers from DNA in the PPS region^2,3,20^. One hallmark of primed adaptation is that nearly all PPS-region sequences from which spacers are acquired contain a consensus 5’-AAG-3’/3’-TTC-5’ PAM (PAM^AAG^)^2,3,20^. A second hallmark of primed adaptation is that spacer acquisition occurs in a bidirectional, orientation-dependent manner relative to the PAM of the PPS. In particular, the non-transcribed strand of spacers acquired from the PAM-proximal region (upstream) or PAM-distal region (downstream) is derived from the nontarget strand or target strand, respectively^21^.

Available *in vivo* models of primed adaptation that contain a plasmid-borne PPS or phage-borne PPS are limited due to difficulties in detecting bidirectional spacer acquisition or by high rates of cell lysis^2,3,21^. Therefore, to overcome limitations of these systems we constructed a derivative of *E. coli* K12 with a type I-E CRISPR-Cas locus containing a spacer, Sp^*yihN*^ encoding a crRNA targeting a chromosomal protospacer in the non-essential gene *yihN* (Fig. 1a; Supplementary Table 1). Induction of *cas* gene expression in “self-targeting” cells leads to inhibition of cell growth accompanied by an increase in cell length (Fig. 1b). Furthermore, analysis of chromosomal DNA by high-throughput sequencing shows induction of *cas* gene expression causes a dramatic loss of ~300 kb of chromosomal DNA in the PPS region (Fig. 1c, Extended Data Fig. 1a-1b. Supplementary Table 2). Loss of PPS-region DNA is also observed in cells containing a catalytically inactive Cas 1 variant (Cas1^H208A^)^22^ but is not observed in cells containing a nuclease-deficient Cas3 variant (Cas3^H74A^)^11^ or cells in which Sp^*yihN*^ is replaced by a spacer targeting M13 phage (Sp^M13^)^10^ (Extended Data Fig. 1a. Supplementary Table 3). Similar results are obtained using methods for analysis of doublestranded or single-stranded DNA (Extended Data Fig. 1b, Supplementary Table 2), indicating that interference-driven degradation of both the target and nontarget strands occurs in the self-targeting strain. The results establish that induction of *cas* gene expression results in interference-driven degradation of PPS-region DNA in the type I-E CRISPR-Cas self-targeting system.

**Figure 1.**
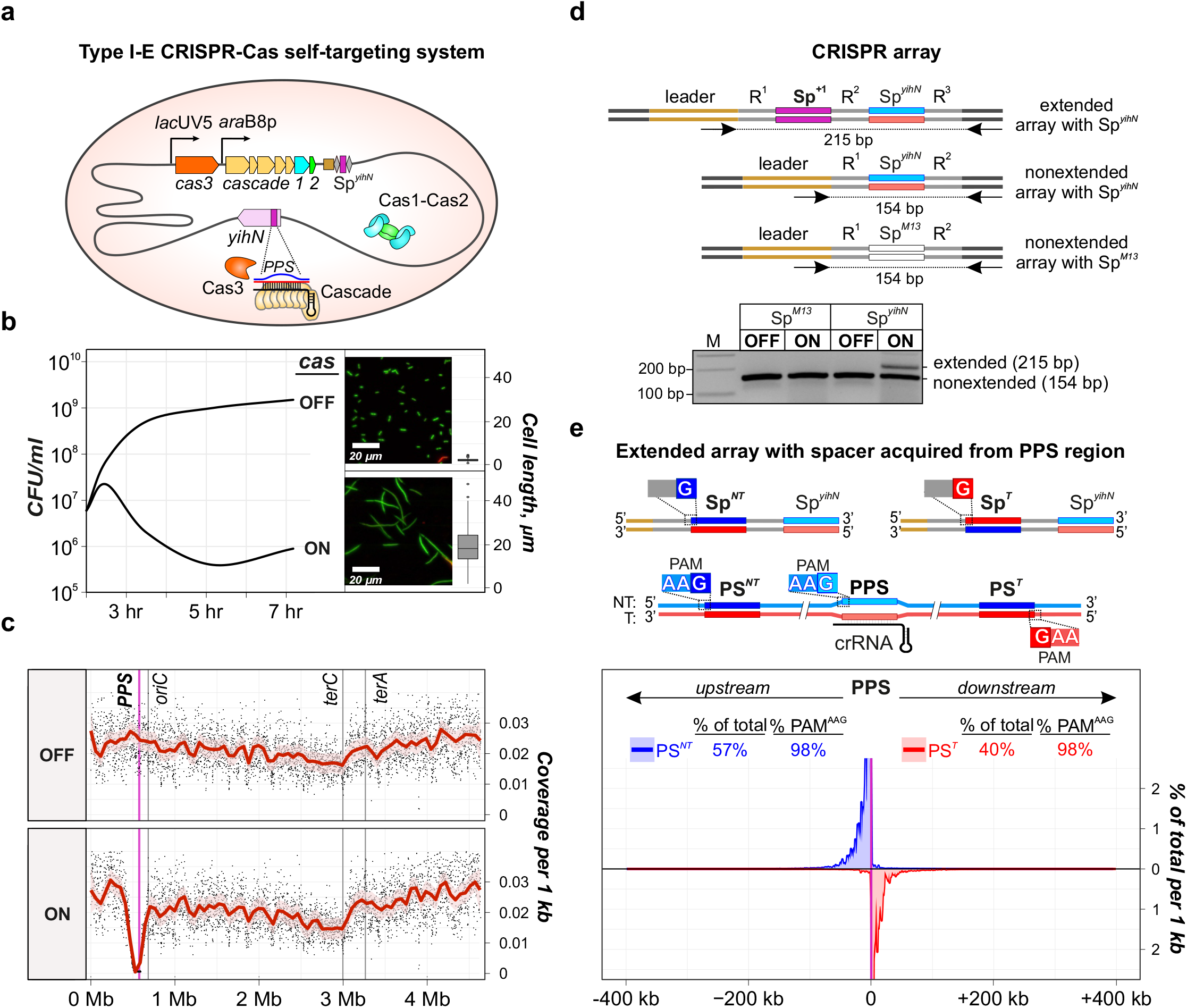
Interference-driven DNA degradation coupled with spacer acquisition in a type I-E CRISPR-Cas self-targeting model system. **a**, Type I-E CRISPR-Cas self-targeting system. Shaded oval, *E. coli* cell; grey line, chromosome; orange, tan. blue, and green pentagons, *cas* genes; brown rectangle, CRISPR-array leader sequence; grey diamonds, repeat sequences; purple rectangle, spacer targeting *yihN* (Sp^*yihN*^); /*ac*UV5 and *ara*B8p, inducible promoters; mauve pentagon, *yihN*: CasS and Cascade, interference machinery; Cas1-Cas2, adaptation machinery; PPS, priming protospacer within *yihN;* blue line, nontarget strand; red line, target strand; black line, crRNA. **b,** Effect of self-targeting on cell growth. Growth curve for cultures in which *cas* gene expression is induced (ON) or not induced (OFF) at time 0. Cells from cultures at 5 hr were used for images (green, viable cells; red, non-viable cells) and boxplot (n = 125; whiskers represent 1.5 × IQR). **c**, Effect of self-targeting on genomic DNA content. *oriC*, site of replication origin; *terA* and *terC*, sites of replication termination; dot, coverage per 1 kb; red line, Loess smoothing; pinkshading, 99% confidence interval, **d**, Effect of self-targeting on spacer acquisition: PCR analysis. Top, extended and nonextended arrays and PCR amplicons. Bottom, results. Sp^+1^, acquired spacer; blue line, non-transcribed strand of Sp^*yihN*^; red line, transcribed strand of Sp^*yihN*^ (directs synthesis of crRNA); R, repeat sequence. M, double-stranded DNA marker, **e**, Effect of self-targeting on spacer acquisition: high-throughput sequencing analysis. Top, extended arrays with spacers acquired from PPS-region protospacers. Bottom, results. Sp^*NT*^, spacer with non-transcribed strand derived from nontarget strand (NT, blue) and transcribed strand derived from target strand (T, red); Sp^*T*^, spacer with non-transcribed strand derived from target strand (T, red) and transcribed strand derived from nontarget strand (NT, blue). PS^*NT*^ protospacer for Sp^*T*^; PS^*T*^, protospacer for Sp^*T*^. Plot shows percentage of spacers per 1 kb derived from PS^*NT*^ (blue) or PS^*T*^ (red). Mean of three biological replicates is shown.

To determine whether interference-driven degradation of PPS-region DNA is coupled with spacer acquisition from PPS-region sequences we analyzed CRISPR arrays by PCR (Fig. 1d). Results indicate ~20% of arrays acquire a spacer in cells in which *cas* gene expression is induced, while no spacer acquisition is detected in cells in which *cas* gene expression is not induced (Fig. 1d). Furthermore, no spacer acquisition is detected in cells in which Sp-^*yith*^ is replaced by Sp^*M13*^ (Fig. 1d), indicating that spacer acquisition requires interference-driven degradation of PPS-region DNA. Iiigh-throughput sequencing analysis of amplicons derived from arrays that have acquired a spacer indicate the self-targeting system exhibits the defining hallmarks of primed adaptation. In particular, >95% of spacers are acquired from a PAM^AAG^-containing “protospacer” in the PPS region and, furthermore, spacer acquisition occurs in a bidirectional, orientation-dependent manner characteristic of the *E. coli* I-E system^21^ (Fig. 1e, Supplementary Table 4–5). We conclude that the type I-E CRISPR-Cas self-targeting strain provides a robust *in vivo* model system for primed adaptation.

It has been proposed that interference-driven DNA degradation produces fragments that serve as spacer precursors in primed adaptation^3,23^. To test this model, we developed a method for strand-specific, high-throughput-sequencing of DNA fragments, “FragSeq,” and applied this method to identify products of degradation in self-targeting cells undergoing primed adaptation (Fig. 2a, Extended Data Fig. 2–4, Supplementary Tables 6–12 and Methods). Results show accumulation of fragments derived from PPS-region DNA in wild-type cells but not in cells containing inactive variants of Cas 1 or Cas3, or cells in which Sp^*yihN*^ is replaced by Sp^*M13*^ (Fig. 2a, Extended Data Fig. 3a, Supplementary Table 7). Thus, accumulation of PPS-region-derived fragments in cells undergoing primed adaptation requires the functional integrity of both interference and adaptation.

**Figure 2.**
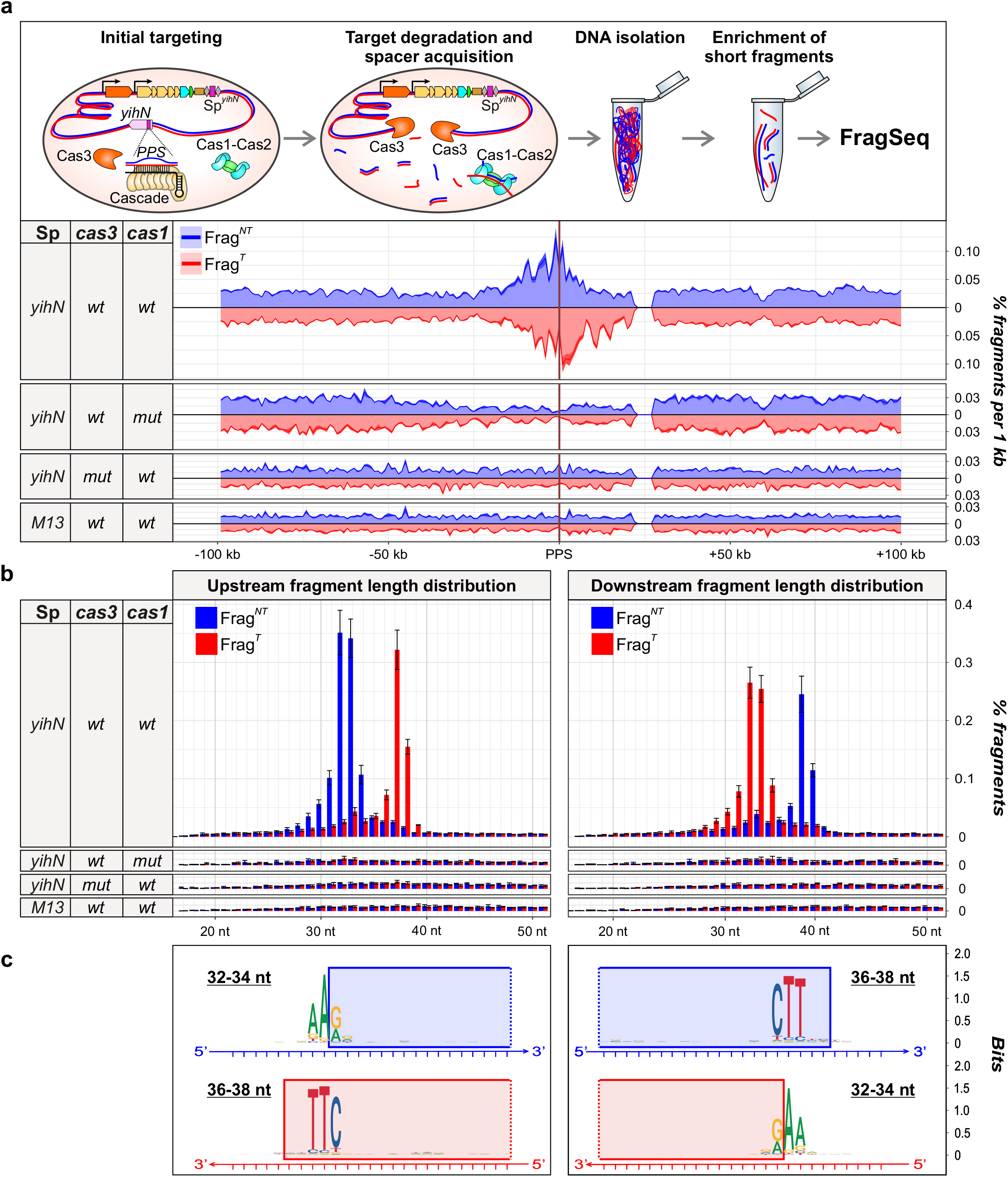
Detection of DNA fragments generated in cells undergoing primed adaptation in the type I-E self-targeting system by FragSeq. **a**, Effect of seif-targeting on PPS-region DNA fragment distributions. Top, events occurring in cells upon induction of *cas* gene expression. Bottom, FragSeq results. Coverage plots show mean of three biological replicates. Blue, nontarget-strand-derived fragments (Frag^*NT*^); red, target-strand-derived fragments (Frag^*T*^). **b**, Length distributions of PPS-region-derived fragments (mean±SEM of three biological replicates). **c**, Sequence alignments of genomic DNA from which PPS-region fragments are derived. Blue rectangles, sequences present in Frag^*NT*^; red rectangles, sequences present in Frag^*T*^.

Analysis of length distributions of the PPS-region-derived fragments indicate they are produced in a bidirectional, orientation-dependent manner reminiscent of spacer acquisition (Fig. 2b). The most abundant nontarget-strand fragments (Frag^NT^) and target-strand fragments (Frag^T^) emanating from the PAM-proximal region of the PPS (upstream) are 32- to 34-nt and 36- to 38-nt, respectively, and the most abundant Frag^NT^ and Frag^1^ emanating from the PAM-distal region of the PPS (downstream) are 36- to 38-nt and 32- to 34-nt, respectively (Fig. 2b). In addition, the relative abundance of complementary 32- to 34-nt and 36- to 38-nt fragments show a positive correlation (Pearson correlation coefficient 0.48, Supplemental Table 11), suggesting the fragments identified by FragSeq represent individual strands of double-stranded DNA products having lengths similar to that of spacers (~30-bp). Alignments of the chromosomal sequences associated with the 5’ or 3’ ends of complementary fragments reveals the presence of a consensus 5’-AAG-3’/3’-TTC-5’ PAM derived from sequences associated with the 5’-ends of 32- to 34-nt fragments and the 3’-ends of 36- to 38-nt fragments (Fig. 2c, Supplementary Tables 9,10). Thus, the results of FragSeq suggest cells undergoing primed adaptation accumulate 33- or 34-bp double-stranded DNA fragments containing a 3’-end, 4- or 3-nt overhang derived from excision of a PAM-containing sequence (Fig. 2c). Furthermore, the relative abundance of these fragments and spacers acquired during primed adaptation that have an identical sequence shows a positive correlation (Pearson correlation coefficient 0.5-0.6, Supplemental Table 12). Accordingly, the results strongly suggest the fragments accumulating in cells undergoing primed adaptation are products of an intermediate step between protospacer selection and spacer integration.

To directly test whether the PPS-region-derived fragments detected by FragSeq serve as substrates for spacer integration we performed a “prespacer efficiency assay^24^” (Fig. 3a). We tested synthetic mimics corresponding to the most abundant PPS-region-derived fragments (Fig. 3b, Supplementary Table 13). Results show that 33- or 34-bp synthetic mimics containing a 3’-end, 4- or 3-nt overhang on the PAM-derived end, respectively, and a blunt PAM-distal end were integrated into arrays with an efficiency similar to a control fragment containing a consensus PAM^AAG^ (~10% prespacer efficiency; Fig. 3b, Supplementary Tables 14, 15). In addition, the synthetic mimics and PAM^AAG^-containing control fragment were integrated in a “direct” orientation with the G:C of the PAM positioned adjacent to the first repeat in the array (Fig. 3, Supplementary Table 15). Introduction of a 5’-end, 1-nt overhang on the PAM-distal end reduced prespacer efficiency by ~45-fold, to a value similar to that of a 33-bp control fragment without a PAM (Fig. 3b, Supplementary Table 15). The results establish that PPS-region-derived fragments containing a 3’-end overhang on the PAM-derived end and blunt PAM-distal end function as efficient spacer precursors.

**Figure 3.**
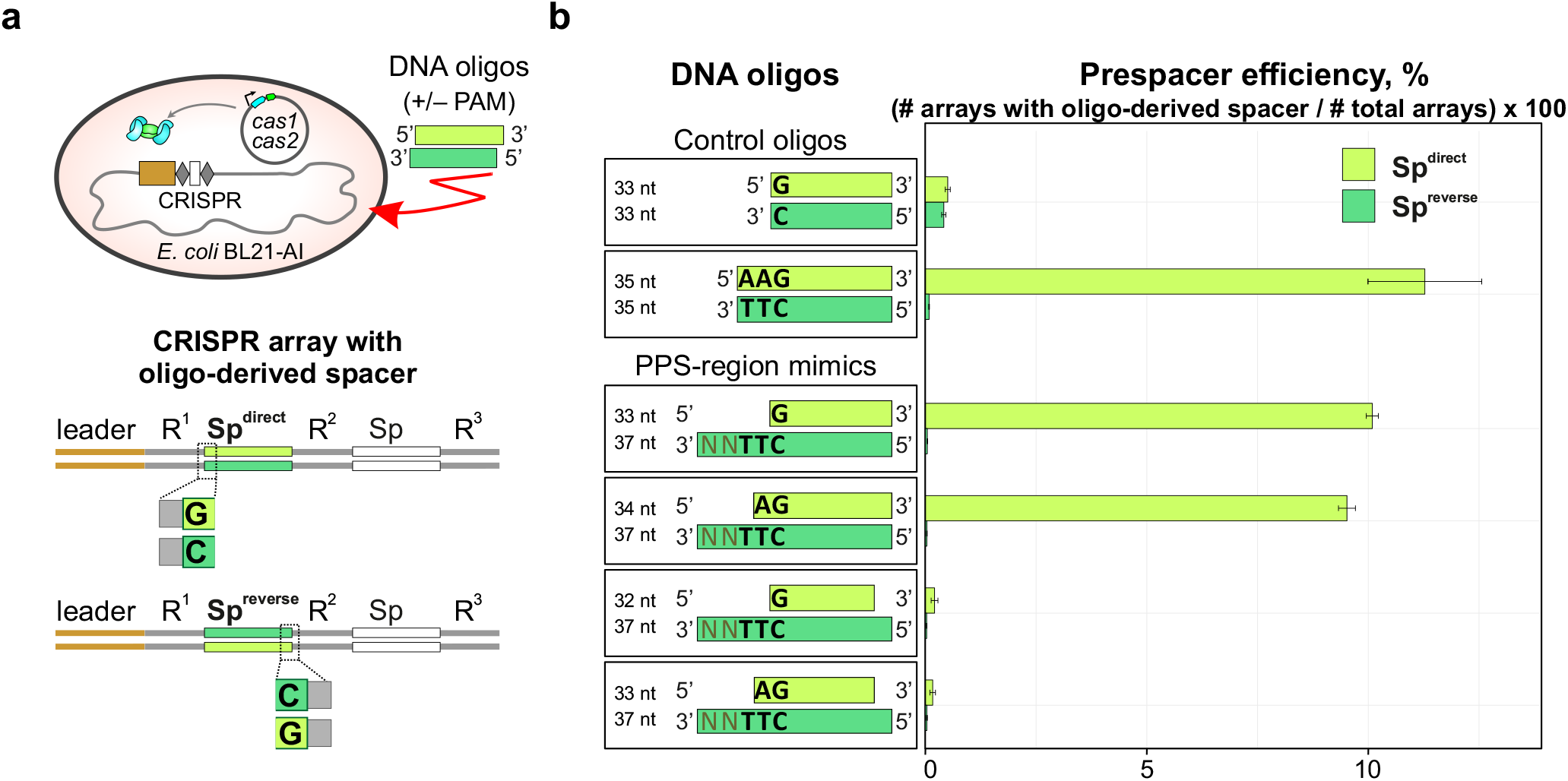
Synthetic mimics of DNA fragments detected in cells undergoing primed adaptation function as spacer precursors. **a**, Prespacer efficiency assay. Top, introduction of synthetic DNA into cells containing a CRISPR array and plasmid that directs expression of *cas1* and *cas2*. Bottom, integration of synthetic DNA into the CRISPR array occurs in either a direct (Sp^direct^) or reverse (Sp^reverse^) orientation, **b**, Results. Left, oligonucleotides analyzed. Right, percentage of arrays containing oligo-derived spacers having a direct (light green) or reverse (dark green) orientation (mean ±SEM of three biological replicates).

In prior work we developed an *E. coli* strain that provides a model system for studies of self-targeting by the type I-F CRISPR-Cas system from *Pseudomonas aeruginosa*^25^ (Extended Data Fig. 5a). Compared with the orientation bias in spacer acquisition observed in type I-E systems, orientation bias in type I-F systems is reversed. In particular, the nontranscribed strand of spacers acquired from the PAM-proximal region of the PPS (upstream) or PAM-distal region of the PPS (downstream) are derived from the target strand or nontarget strand, respectively in type I-F. To determine whether spacer precursors could be detected in the type I-F system we performed FragSeq analysis in cells undergoing primed adaptation (Extended Data Fig. 5b, Supplementary Tables 17–21). Similar to the type I-E system, we detect accumulation of spacer-sized double-stranded DNA fragments containing a 3’-end. 5-nt overhang on the PAM-derived end and a blunt PAM-distal end (Extended Data Fig. 5b). Thus, in spite of exhibiting opposite orientation bias in spacer acquisition, primed adaptation in type I-E and type I-F systems involves generation of spacer precursors with a similar structure (Extended Data Fig. 5c).

In summary, we have identified spacer precursors produced as products of an intermediate step (or steps) between protospacer selection and spacer integration for type I-E and type I-F CRISPR-Cas systems. Accumulation of spacer precursors in the type I-E system requires the functional integrity of components of interference and adaptation (Fig. 4) indicating that protospacer selection involves coordination between the interference machinery and adaptation machinery (Fig. 4a). Strikingly, spacer precursors detected during primed adaptation in both type I-E and type I-F systems share an asymmetrical structure characterized by a 3’-end overhang on the PAM-derived end. Thus, we propose that spacer precursors detected in this work are products generated during universal steps of prespacer processing that occur in all type I CRISPR-Cas systems. We further propose that the asymmetrical structure of the spacer precursors detected in this work is a key determinant of the sequential integration of prespacers into the CRISPR array (Fig. 4b). In addition, the FragSeq method reported in this work should be applicable, essentially without modification, to identify spacer precursors that form *in vivo* in any CRISPR-Cas system.

**Figure 4.**
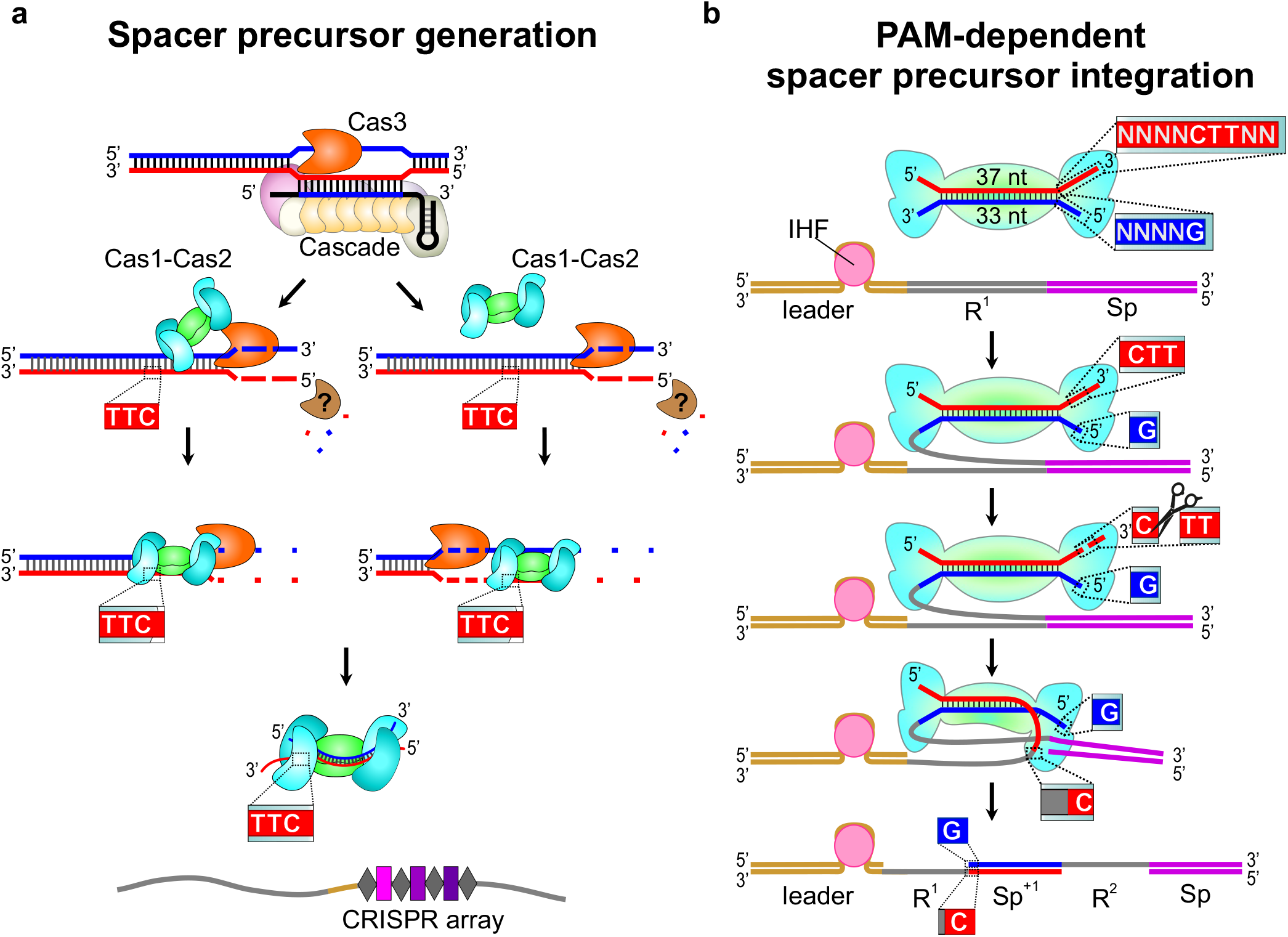
Model of primed adaptation in type I-E CRISPR-Cas systems. **a**, Generation of spacer precursors involves coordination between interference and adaptation. Pathway on left depicts “direct” coordination between interference and adaptation in which Cas1-Cas2 associates with Cas3 as it moves along DNA^16,26^. Pathway on right depicts “indirect” coordination between interference and adaptation in which Cas1-Cas2 captures products of Cas3-mediated DNA degradation^3,23^. Both pathways generate spacer precursors containing a 3’-end overhang on the PAM-derived end and blunt PAM-distal end. Model depicts rapid degradation of DNA not selected as a spacer precursor by Cas3 and an unknown nuclease (brown), **b**, Sequential integration of spacer precursors. First, binding of IHF to the leader stimulates integration of the blunt PAM-distal end between the leader and first repeat sequence^27^. Second, Cas1-mediated cleavage of the 3’-overhang present on the PAM-derived end facilitates integration between the first repeat sequence and first spacer of the array^28^. The order of events depicted results in integration of the spacer precursor in a direct orientation with respect to the PAM (see Fig. 3a).

## Acknowledgments

The work was carried out using scientific equipment of the Center of Shared Usage «The analytical center of nano- and biotechnologies of SPbPU and Next Generation Sequencing equipment of The Waksman Genomics Core Facility. This work was supported by NIH grant GM10407 (KS), NIH grant GM118059 (BEN), and Russian Science Foundation grant 14-14-00988 (KS).

## Contributions

A. A. S, I. O. V., B. E. N., K. S. and E. Semenova designed the experiments.

A. A. S., E. Savitskaya, K. A. D., I. O. V., I. F., N. M., A. M., A. S. and E. Semenova performed the experiments.

A. A. S. and E. Savitskaya analyzed high-throughput sequencing data.

A. A. S., B. E. N., K. S. and E. Semenova wrote the manuscript.

## Competing Interests

The authors declare no competing interests

## Methods

### Bacterial strains and plasmids

The *E. coli* strains used in this study are listed in Supplementary Table 1. Red recombinase-mediated gene-replacement technique was used to obtain strains KD403, KD518 and KD753^29^.

Plasmid pCDF-Ec.cas1/2 was constructed as described previously^4^. Plasmids pCas and pCsy were described earlier^25^.

### Growth conditions during CRISPR interference and primed adaptation assays

For analysis of CRISPR-mediated self-targeting by the type I-E system, overnight culture of KD403 strain grown at 37°C in LB medium was diluted 100-fold into 10 ml fresh LB and incubated at 37°C until OD_600_ reached 0.3. Culture was divided into two portions, *cas* genes inducers, IPTG and L-(+)-arabinose, were added at 1 mM concentration to one portion, and cultures with and without inducers were incubated at 37°C for 7 hr. At various time points postinduction, cells were plated with serial dilutions on 1.5% LB agar plates for counting colony forming units (CFU) or were monitored using fluorescent microscopy.

In assays using strains KD403, KD518, KD753 and KD263 that were followed by sequencing of total genomic DNA, short DNA fragments or newly acquired spacers, similar conditions of culture growth and *cas* genes induction were applied, except that overnight cultures were diluted 100-fold in 100 ml LB and grown at 30°C. Five hours postinduction, 10 ml of cells were pelleted by centrifugation at 3000 × *g* for 5 min at 4°C, washed with 10 ml of PBS, pelleted by centrifugation at 3000 × *g* for 5 min at 4°C and resuspended in 1 ml of PBS. Cells were divided into 125 μl aliquots and stored at −70°C before they were used for DNA isolation.

For analysis of short DNA fragments generated during self-targeting by the type I-F system, cultures of strain KD675 transformed with plasmids pCas and pCsy were grown at 37°C in LB supplemented with 100 μg/ml ampicillin and 50 μg/ml spectinomycin. Overnight cultures were diluted 200-fold into 10 ml of LB without antibiotics, grown at 37°C until OD_600_ reached 0.3 and supplemented with 1mM IPTG and 1mM L-(+)-arabinose. Cells were harvested 24 hr postinduction and prepared for DNA isolation as described above for strains KD403, KD518, KD753 and KD263.

### Fluorescence microscopy

Cultures grown with or without induction of *cas* gene expression were analyzed using a LIVE/DEAD viability kit (Thermo Scientific) at 5 hr after induction. Viable cells in each culture were detected by addition of 20 μM SYT09, green fluorescent dye that can penetrate through intact cell membranes. Non-viable cells in each culture were detected by addition of 20 μM propidium iodide dye, which cannot enter viable cells. Sample chambers were made using a microscope slide (Menzel-Gläser) with two strips on the upper and lower edges formed by double-sided sticky tape (Scotch TM). To obtain a flat substrate required for high-quality visualization of bacteria, a 1.5% agarose solution was placed between tape strips and covered with another microscopic slide. After solidification of the agarose the upper slide was removed and several agarose pads were formed. 1 μl of each cell suspension (with and without induction) was placed on an agarose pad. The microscopic chamber was sealed using coverslip (24 × 24 mm, Menzel-Gläser).

Fluorescence microscopy was performed using Zeiss AxioImager.Z1 upright microscope. Fluorescence signals in green (living cells) and red (dead cells) fluorescent channels were detected using Zeiss Filter Set 10 and Semrock mCherry-40LP filter set respectively. Fluorescent images of self-targeting cells were obtained using Cascade II:1024 back-illuminated EMCCD camera (Photometrics). Microscope was controlled using AxioVision Microscopy Software (Zeiss). All image analysis was performed using ImageJ (Fiji) with ObjectJ plugin used for measurements of cell length^30^.

### High-throughput sequencing of total genomic DNA

Total genomic DNA was purified by GeneJET Genomic DNA Purification Kit (ThermoFisher Scientifc). Sequencing libraries were prepared either by NEBNext^®^ Ultra™ II DNA Library Prep Kit for Illumina (NEB) or by Accel-NGS^®^ 1S Plus DNA Library Kit (Swift Biosciences) and sequenced on a NextSeq 500 platform.

Raw reads were analyzed in R with ShortRead and Biostrings packages^31^. Bases with quality < 20 were substituted with N and the reads were mapped to the KD403 reference genome using Unipro UGENE platform^32^. Bowtie2 was used as a tool for alignment with end-to-end alignment mode and 1 mismatch allowed^33^. The BAM files were analyzed by Rsamtools package and reads with the MAPQ score equal to 42 were selected and used for downstream coverage analysis^34^. Mean coverage over nonoverlapping 1 kb bins was calculated and normalized to the total coverage (the sum of means).

### High-throughput sequencing of spacers acquired during primed adaptation

Cell lysates were prepared by resuspending cells in water and heating at 95°C for 5 min. Cell debris was removed from lysates by centrifugation at 16 × g for 1 min. For analysis of spacer acquisition in strains KD263 and KD403 lysates were used in PCR reactions containing primers LDR-F2 (ATGCTTTAAGAACAAATGTATACTTTTAG) and Ec.min_R (CGAAGGCGTCTTGATGGGTTTG) (25 cycles, T_a_ = 52°C). Reaction products were separated by agarose gel electrophoresis (Fig. 1d). To obtain amplicons derived from extended CRISPR arrays in strain KD403 PCR reactions were performed using primers LDR-F2 (ATGCTTTAAGAACAAATGTATACTTTTAG) and autoSP2_R (AATAGCGAACAACAAGGTCGGTTG) (30 cycles, T_a_=52°C). Reaction products were separated by agarose gel electrophoresis and the amplicon derived from the extended array was purified from the gel using a GeneJET Extraction Kit (ThermoFisher Scientifc) and sequenced on a NextSeq 500 system.

Bioinformatic analysis was performed in R using ShortRead and Biostrings packages^31^. Bases with quality < 20 were substituted with N and spacer sequences were extracted from the reads containing two or more CRISPR repeats. Spacers of length 33 bp were mapped to the KD403 genome to identify 33-bp “protospacer” sequences with 0 mismatches. Spacers that aligned to one position in the chromosome were used to determine protospacer distribution along the genome. Spacers arising from protospacers due to potential ‘slippage’ or ‘flippage’ were removed from analysis^35^ (Supplementary Tables 4, 5).

### Prespacer efficiency assay

Competent cells were prepared from a culture of BL21-AI cells containing plasmid pCDF-Ec.cas1/2 grown in the presence of 13 mM arabinose and 1 mM IPTG to induce expression of *cas1* and *cas2*. Complementary oligonucleotides (Supplementary Table 13) were introduced into cells by electroporation as described previously^24^. Lysates from cell cultures were generated and used in PCR reactions containing a primer complementary to the leader sequence (GGTAGATTGTGACTGGCTTAAAAAATC) and a primer complementary to the preexisting spacer in the array (GTTTGAGCGATGATATTTGTGCTC), respectively. Amplicons corresponding to extended and nonextended CRISPR arrays were isolated using GeneJET PCR Purification Kit (ThermoFisher Scientifc) and sequenced on a NextSeq 500 platform. Bioinformatic analysis was performed in R using ShortRead and Biostrings packages^31^. Bases with quality < 20 were substituted with N and reads containing the leader-repeat junction were further analyzed (sequence spanning the last 21 nucleotides of the leader and the whole repeat were searched for with 6 mismatches allowed). The reads containing two repeats were considered expanded. Newly acquired spacers were extracted from the expanded reads and mapped to the genome, plasmid and transforming oligonucleotide sequence with 2 mismatches allowed. 33 bp oligo-derived spacers that were cut between AA and G before integration were considered as properly processed. For simplicity only properly processed oligo-derived spacers inserted into the CRISPR array in direct (GCCCAATTTACTACTCGTTCTGGTGTTTCTCGT) or reverse (ACGAGAAACACCAGAACGAGTAGTAAATTGGGC) orientation were included into analysis.

### Isolation of DNA fragments generated *in vivo*

Total genomic DNA was isolated from cultures of strains KD403, KD518, KD753, KD263 and KD675 by collecting 1.25 ml of cell suspensions by centrifugation, resuspending cells in 125 μl of PBS, adding 2 ml lysis buffer (0.6% SDS, 12 μg/ml proteinase K in lx TE buffer) and incubating at 55°C for 1 hr. Two milliliters of phenol:chloroform:isoamyl alcohol (25:24:1) (pH 8) were added to the lysate, the solution was gently mixed, and the aqueous and organic phases separated by centrifugation at 7000 × *g* for 10 min at room temperature. The upper aqueous phase containing total genomic DNA was collected and the residual phenol was removed by the addition of 2 ml of chloroform:isoamyl alcohol (24:1). The solution was gently mixed, centrifuged at 7000 × *g* for 10 min at room temperature, the upper DNA-containing fraction was transferred into fresh tube, 0.2 M NaCl, 15 μg/ml of Glycoblue (Invitrogen) and two volumes of cold 100% ethanol was added, and the solution was incubated at −80°C overnight. Precipitated DNA was recovered by centrifugation at 21000 × *g* for 30 min at 4°C. Pellets were washed twice with 80% ethanol, resuspended in 200 μl of 1x TE buffer, and treated with 1 mg/ml RNase A at 37°C for 30 min to remove residual RNA. DNA was isolated by phenol:chloroform:isoamyl alcohol extraction and ethanol precipitation as described above.

DNA fragments < 700 bp in length were isolated from 9 μg of total genomic DNA using a Select-a-Size DNA Clean & Concentrator kit (Zymo Research) according to manufacturer’s recommendations. To ensure the binding of fragments <50 bp to the column filter, the volume of 100% ethanol added to the fraction prior to on-filter purification was increased from 290 μl to 600 μl. DNA fragments were eluted with 2 × 50 μl of elution buffer, pooled aid purified by ethanol precipitation. 100 μl DNA was mixed with 10 μl of 3 M NaOAc (O.lxV), 1 μl of 10 mg/ml glycogen (0.01xV), and 330 μl of 100% ethanol, vortexed, and incubated overnight at −80°C. DNA was recovered by centrifugation at 21000 × *g* for 30 min at 4°C. Pellets were washed 3 times with 80% cold ethanol, air dried for ~5 min, and resuspended in 5 μl of nuclease-free water.

### High-throughput sequencing of DNA fragments: FragSeq

The DNA oligo i116 that served as a 3’ adapter was adenylated using 5’ DNA Adenylation Kit (NEB), purified by ethanol precipitation as above and diluted to 10 μM with nuclease-free water.

DNA fragments < 700 bp (in 5 μl water) were heat-denatured at 95°C for 5 min, cooled to 65°C, and mixed with 0.5 μM adenylated oligo i116, 1x NEBuffer 1, 5 mM MnC¾, and 10 pmol of thermostable 5’ App DNA/RNA ligase (NEB) in 10-μl reaction volume. The mixture was incubated at 65°C for 1 hr, heated at 90°C for 3 min, and cooled to 4°C on ice. Ligated products were combined with 1x T4 RNA ligase buffer, 12% PEG 8000, 10 mM DTT, 60 μg/ml BSA, and 10 U of T4 RNA ligase 1 (NEB) in a 25-μl reaction volume. The reaction was incubated at 16°C for 16 hr, 25 μl of 2x loading dye was added, and products were separated by electrophoresis on 10% 7 M urea slab gels (equilibrated and run in 1x TBE buffer). The gel was stained with SYBR Gold nucleic acid gel stain, bands were visualized on a UV transilluminator, and products of —40 to —500 nt were excised from the gel and recovered as described in Vvedenskaya *et al.^36^*. The excised gel slice was crushed, 400 μl of 0.3 M NaCl in 1x TE buffer was added, and the mixture incubated at 70°C for 10 min. The eluate was collected using a Spin-X column. After the first elution step the elution procedure was repeated, eluates were pooled, and DNA was isolated by ethanol precipitation and resuspended in 15 μl of nuclease-free water.

Next, the 3’ adapter-ligated DNA fragments were adenylated using 5’ DNA Adenylation Kit (NEB) in a 20-μl reaction following the manufacturer’s recommendations. Nuclease-free water was added to 100 μl, DNA fragments were purified by ethanol precipitation and resuspended in 5 μl of nuclease-free water. The two-step ligation procedure described above was repeated using 5 μl of adenylated 3’-ligated DNA fragments, 0.5 μM of barcoded oligos i 112, i113, i114 or i115 that served as 5’ adapters (barcodes were used as internal controls; Supplementary Table 22), 10 pmol of thermostable 5’ App DNA/RNA ligase at the first ligation step, and 10 U of T4 RNA ligase 1 at the second ligation step. Reactions were stopped by addition of 25 μl of 2x loading dye, and products were separated by electrophoresis on 10% 7 M urea slab gels (equilibrated and run in 1x TBE buffer). DNA products of ~70 to ~500 nt in size were excised and eluted from the gel as described above, isolated by ethanol precipitation, and resuspended in 20 μl of nuclease-free water.

To amplify DNA, 2 to 8 μl of adapter-ligated DNA fragments were added to a mixture containing 1x Phusion HF reaction buffer, 0.2 inM dNTPs, 0.25 μM Illumina RP1 primer, 0.25 μM Illumina index primer and 0.02 U/μl Phusion HF polymerase in a 30-μl reaction (Supplementary Table 23). PCR was performed with an initial denaturation step of 30 sec at 98°C, amplification for 15 cycles (denaturation for 10 sec at 98°C, annealing for 20 sec at 62°C and extension for 15 sec at 72°C), and a final extension for 5 min at 72°C Amplicons were isolated by electrophoresis using a non-denaturing 10% slab gel (equilibrated and run in 1x TBE). The gel was stained with SYBR Gold nucleic acid gel stain and species of—150 to —300 bp were excised. DNA products were eluted from the gel with 600 μl of 0.3 M NaCl in 1xTE buffer at 37°C for 3 hr, purified by ethanol precipitation, and resuspended in 25 μl of nuclease-free water. Barcoded libraries were sequenced on Illumina NextSeq 500 platform in high output mode.

Bioinformatic analysis was performed in R using ShortRead and Biostrings packages^31^. Bases with quality < 20 were substituted with N. After adapter trimming, all reads were compared to each other to reveal clusters of overamplified reads containing the same insert and combination of unique molecular identifiers conjugated to adapters. For each cluster a consensus sequence was extracted and used together with non-overamplified reads for further alignment to KD403 reference genome with 2 mismatches allowed. Only reads with a length 16 to 100 nt uniquely aligned to the genome were further analyzed (Extended Data Figure 4). Logos were generated using ggseqlogo package^37^.

### Code availability

All codes used in this study are available to editors and reviewers upon request from the corresponding author. The code will be made publicly available at publication.

### Data availability

Raw sequencing data obtained in this study are available to editors and reviewers upon request from the corresponding author. The data will be made publicly available at publication.

---

**Extended Data Fig. 1.**
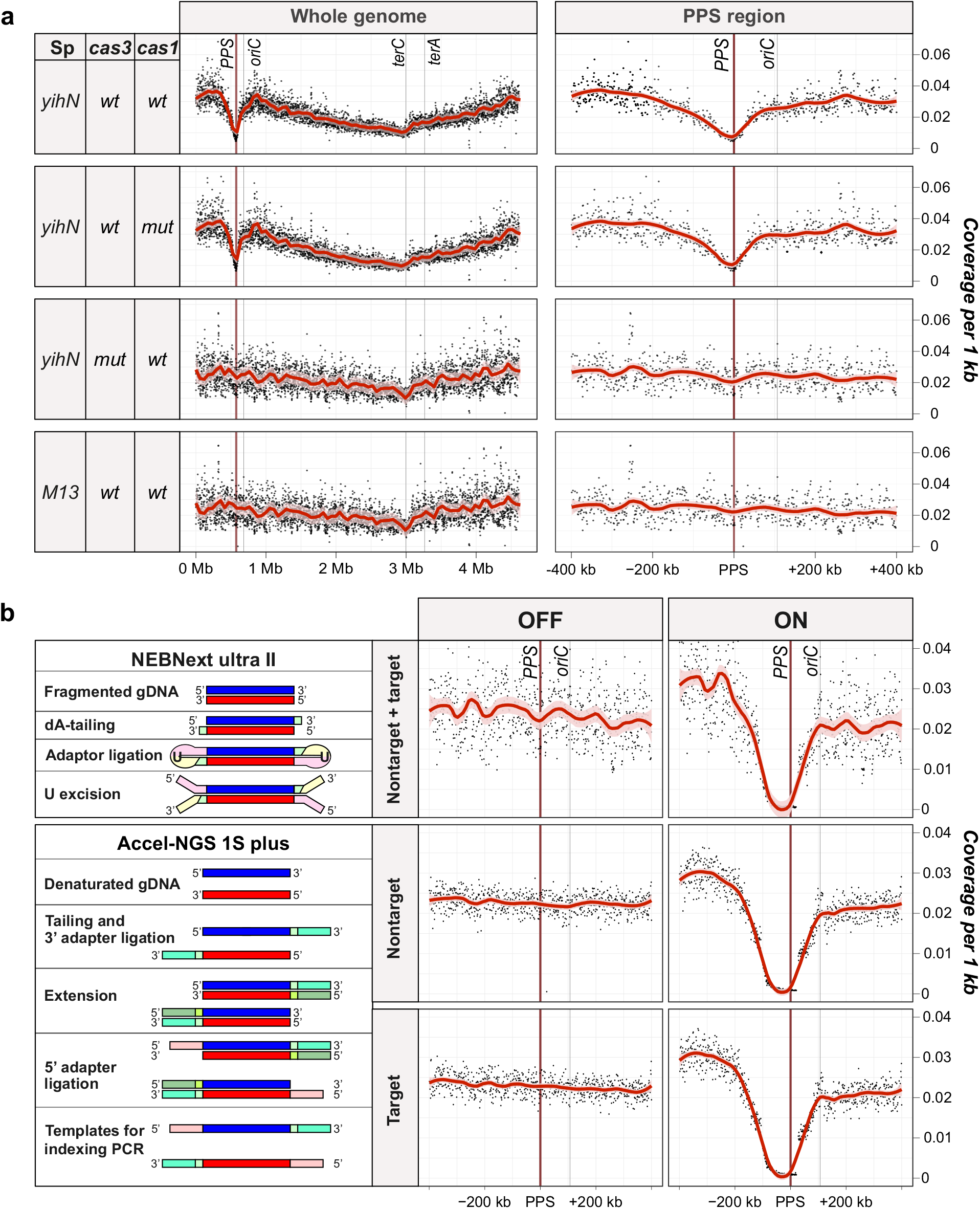
CRISPR interference in the type I-E CRISPR-Cas self-targeting model system results in loss of chromosomal DNA in the PPS^*yihN*^ region. **a,** High-throughput-sequencing analysis of genomic DNA: effects of disruptions in components of interference or adaptation. Graph of sequence coverage per 1 kb for the whole genome (left) or PPS^*yihN*^ region (right) in the indicated strains. *oriC*, site of replication origin; *terA* and *terC*, sites of replication termination; dot, coverage per 1 kb (mean of 3 biological replicates); red line, Loess smoothing; pink shading, 99% confidence interval. *cas1 mut*, gene encoding Cas1^H204A^ *cas3 mut*, gene encoding Cas3^H74A^. (We note that differences in growth rate likely account forthe difference in coverage between *oriC* and the *terC*site in cells undergoing interference vs. cells not undergoing interference; Supplementary Table 3). **b**, High-throughput sequencing analysis of genomic DNA: comparison of library construction methods. Left, steps in library construction using a NEBNext ultra II kit (analysis of double-stranded DNA) or Accel NGS 1S plus kit (analysis of single-stranded DNA). Right, PPS-region coverage plots obtained for wild-type cells.

**Extended Data Fig. 2.**
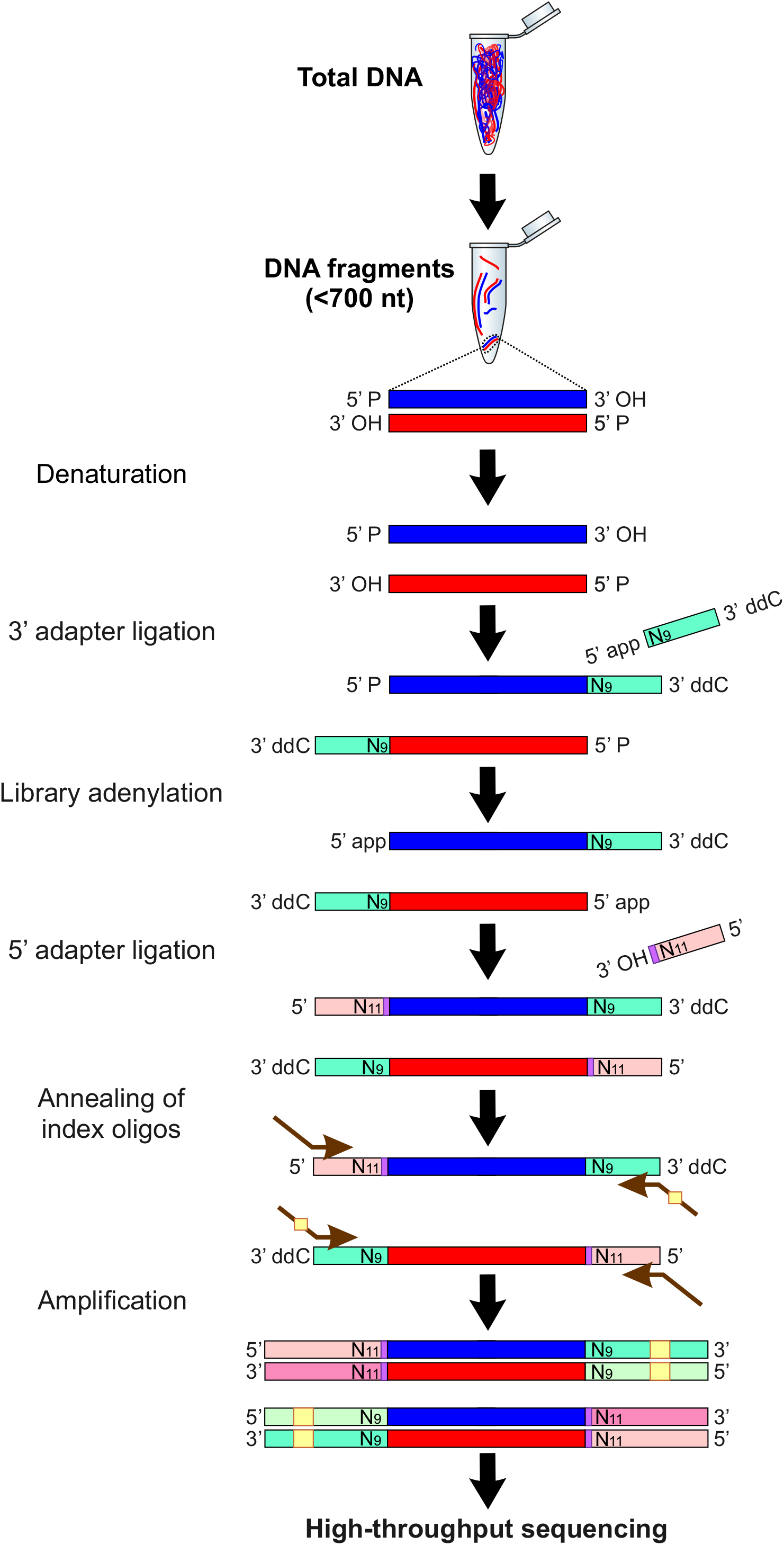
Strand-specific, high-throughput sequencing of DNA fragments, “FragSeq.” Steps in library construction. 5’ app, adenylated 5’ end; 3’ ddC, blocked 3’ end; N9 and N11, unique molecular identifiers on 3’ and 5’ adapters; purple rectangle, 4-n? barcode on 5’ adapter; yellow rectangle, index.

**Extended Data Fig. 3.**
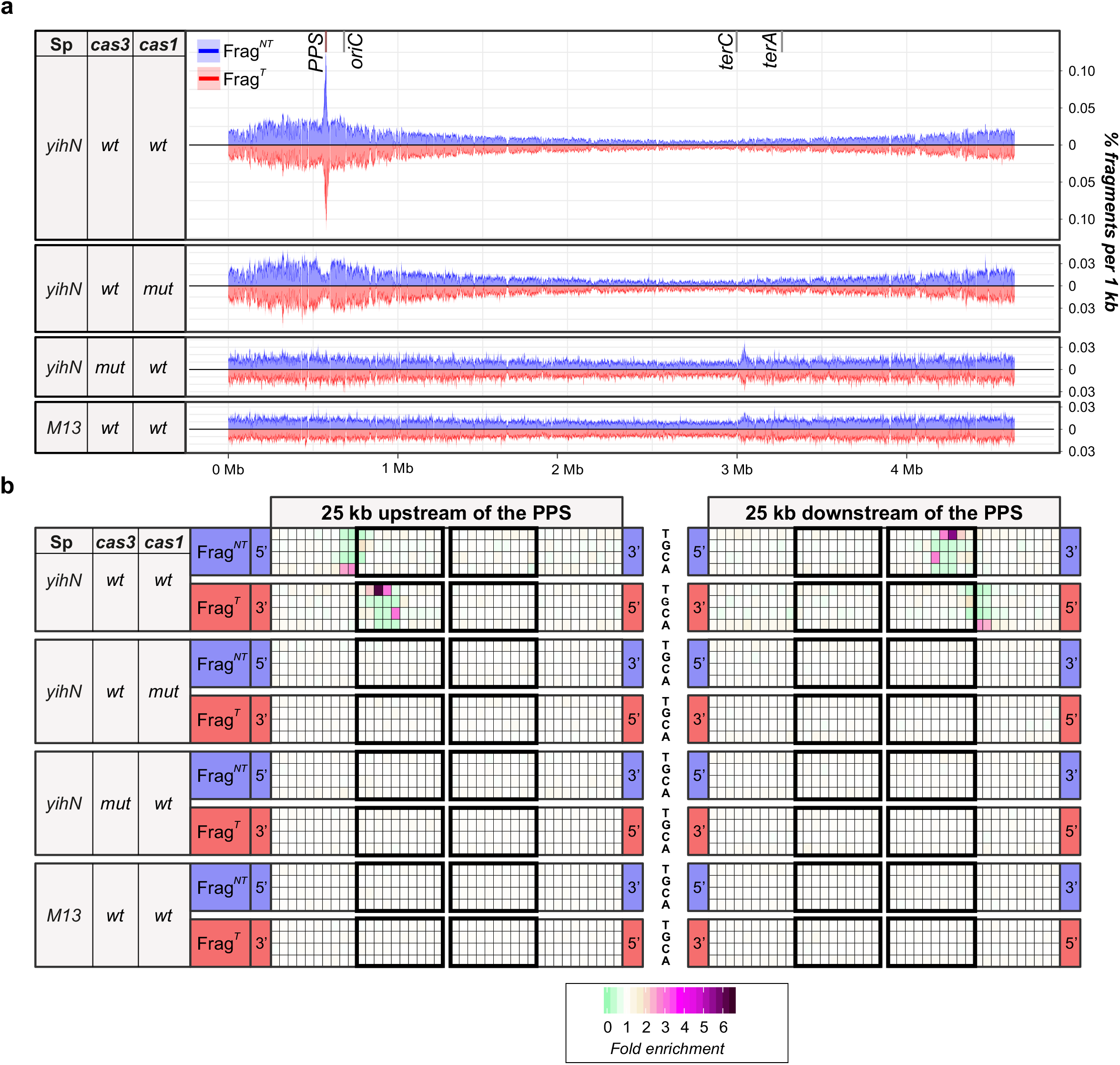
FragSeq results for the type I-E CRISPR-Cas self-targeting model system: fragment coverage plots and sequence analysis. **a**, Genomic coverage plots. Percentage of total DNAfragments per 1 kb for the indicated strains (mean of three biological replicates). Coordinates on the X-axis represent the location on the *E. coli* chromosome. Blue, nontarget-strand-derived fragments (Frag^*NT*^); red, target-strand-derived fragments (Frag^*T*^). **b,** Sequence features of PPS-region fragments and adjacent chromosomal region. Plot shows heat map of relative abundance of A, T, C, or G for the indicated fragment 5’ or 3’ ends. Ten positions of sequences that are detected in fragment 5’ or 3’ ends are shown in black rectangles. Shading represents enrichment (>1) or depletion (< 1) of each nucleotide for sequences associated with PPS-region-derived fragments vs. sequences associated with non-PPS-region-derived fragments.

**Extended Data Fig. 4.**
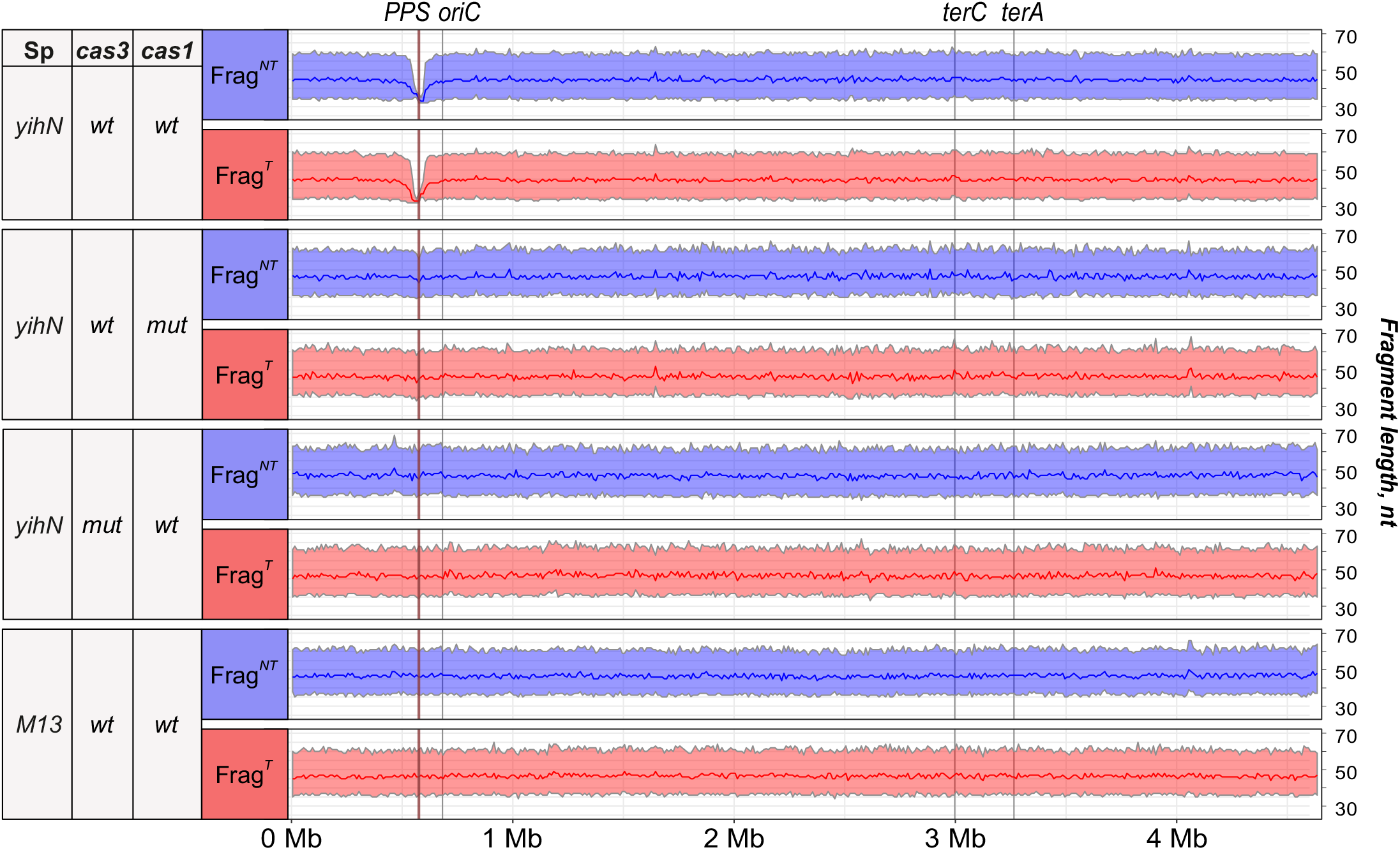
FragSeq results for the type I-E CRISPR-Cas self-targeting model system: length distributions. Length distributions of genome-derived fragments in the indicated strains. Coordinates on the X-axis represent the location on the *E. coli* chromosome. Solid lines represent the median fragment length per 10 kb, shaded areas represent fragment lengths between the first and third quartiles.

**Extended Data Fig. 5.**
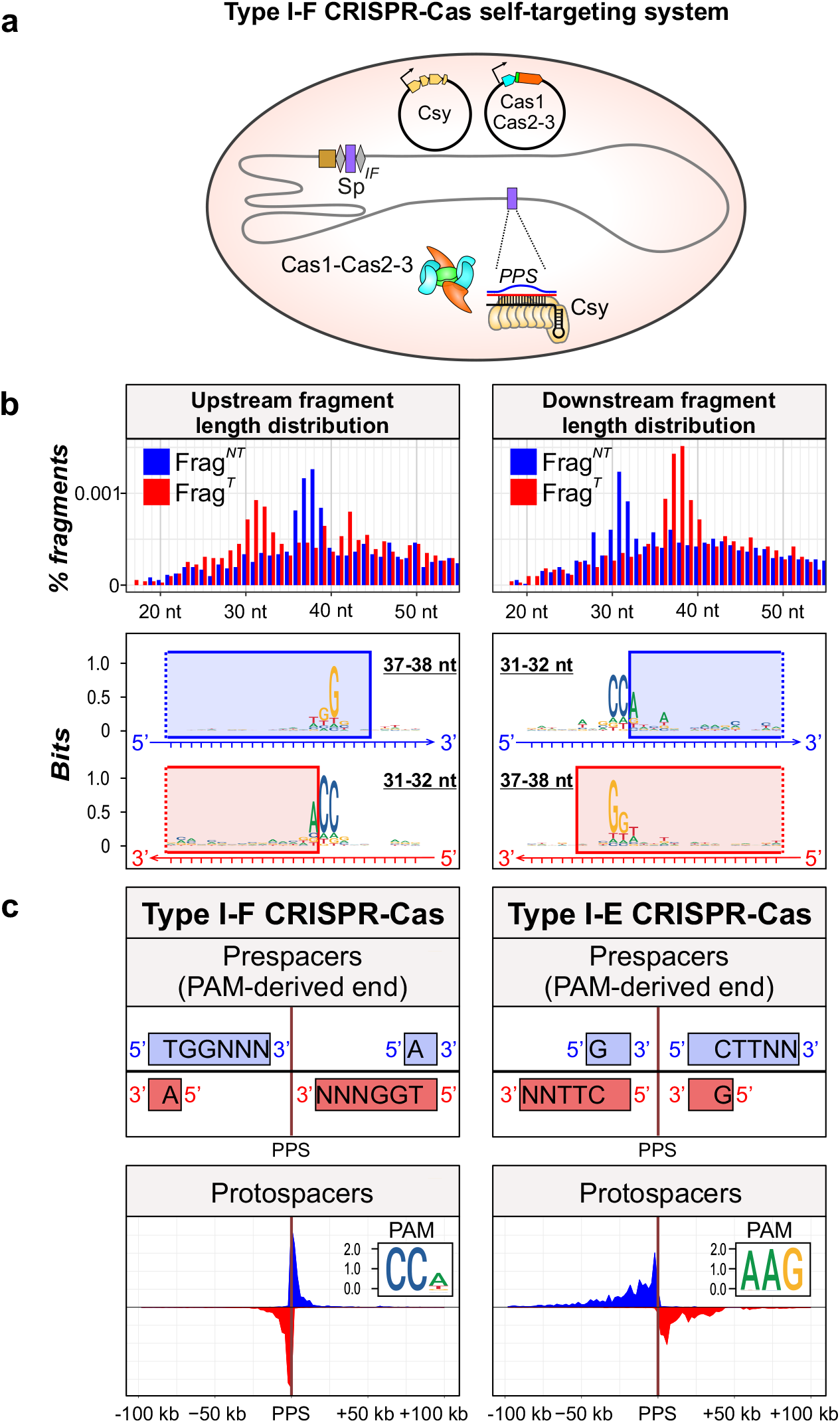
Identification of spacer precursors generated in a type I-F self-targeting system by FragSeq. **a,** Components of type I-F CRISPR-Cas self-interference system. Shaded oval, *E. coli* cell; grey line, chromosomal DNA; black line, plasmid DNA; orange, tan, blue, and green pentagons, cas and csy genes; brown rectangle, array leader sequence; grey diamonds, array repeat sequences; purple rectangles, spacer and chromosomal PPS targeted by spacer-derived crRNA; Csy, type I-F effector complex; Cas1-Cas2-3, complex of Cas1 and Cas2-3 proteins, **b,** FragSeq results: length distributions of fragments (top) and sequence features of PPS-region sequences from which fragments are derived (bottom). Logos for 31 −32-nt fragments were generated by aligning sequences 10-nt upstream to 15-nt downstream of the fragment 5’ end. Logos for 37-38-nt fragments were generated by aligning sequences 20-nt upstream to 5-nt downstream of the fragment 3’ end. Blue rectangles, sequences present in Frag^*NT*^; red rectangles, sequences present in Frag^*T*^ **c**, Comparison of PPS-region-derived fragments and PPS-region protospacers in type I-F and type I-E self-targeting systems. Inset, logo derived from alignment of PPS-region PAMs.

**Supplementary Table 1.**
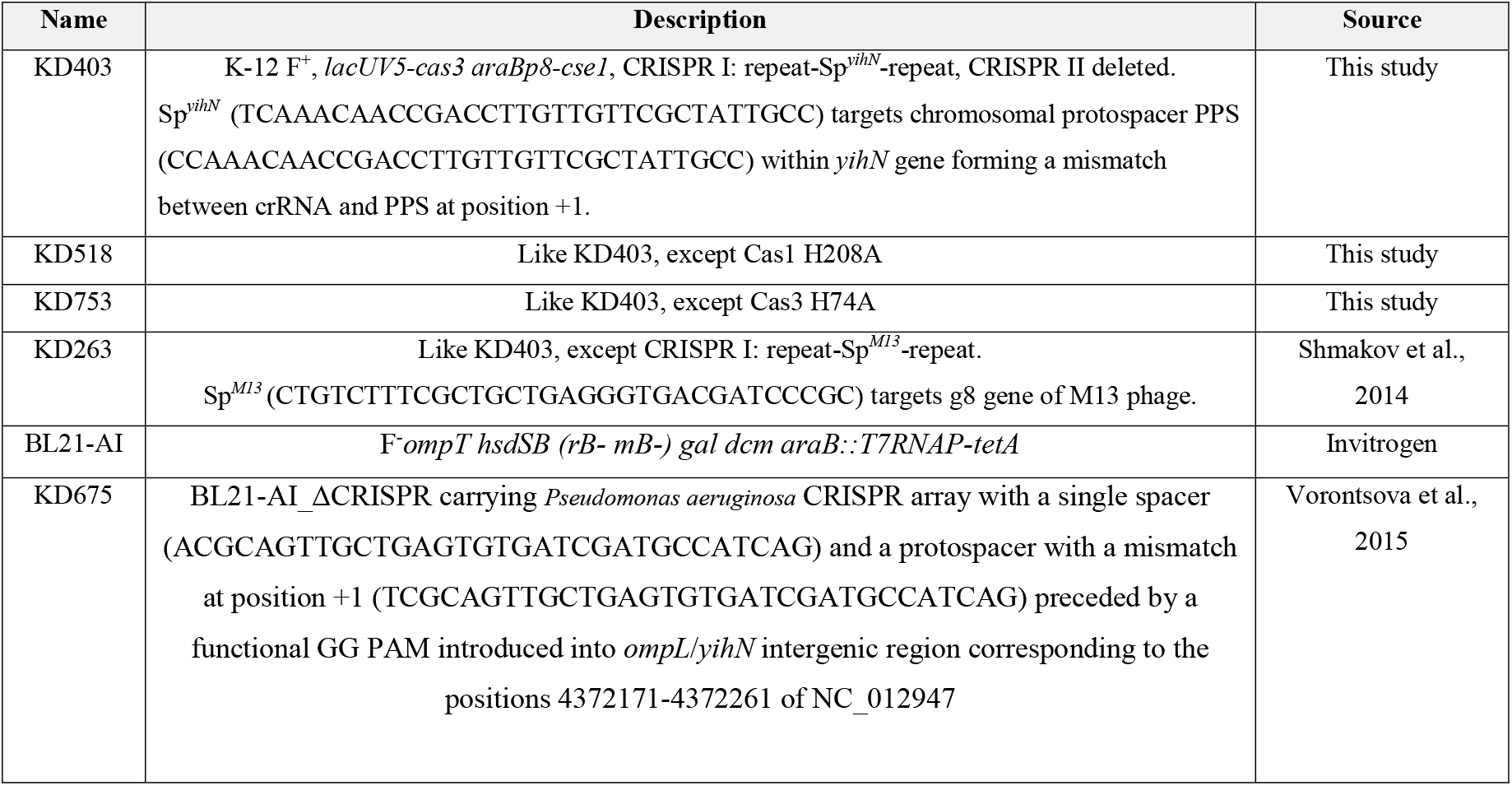
Strains used in this study.

**Supplementary Table 2.** Statistics for sequencing total genomic DNA purified from self-targeting strain KD403 (with or without induction of *cas* genes expression)

| **Library preparation** | **Strain** | **Number of reads aligned to the genome** | **Mean coverage, all genome** | **Mean coverage, PPS-flanking regions\*** | **Mean coverage, PPS** | **Mean coverage, *terC*** | **Ratio of coverage: PPS-flanking regions / PPS** | **Ratio of coverage: PPS-flanking regions / *terC*** |
| --- | --- | --- | --- | --- | --- | --- | --- | --- |
| NEBNext® Ultra™ II DNA Library Prep Kit (NEB) | KD403 -I | 5857376 | 94.8 | 100.7 | 91.5 | 80.3 | 1.1 | 1.3 |
| Accel-NGS® 1S Plus DNA Library Kit (Swift Biosciences) | KD403 -I | 4811616 | 77.8 | 84.9 | 83.7 | 76.9 | 1 | 1.1 |
| NEBNext® Ultra™ II DNA Library Prep Kit (NEB) | KD403 +I | 2890624 | 46.8 | 50.7 | 0.23 | 37.3 | 218.2 | 1.4 |
| Accel-NGS® 1S Plus DNA Library Kit (Swift Biosciences) | KD403 +I | 3061436 | 49.5 | 57.1 | 0.25 | 45.2 | 230.7 | 1.3 |

\*Mean coverage over PPS-flanking regions was calculated as a mean of coverage 200 kb upstream and 100 kb downstream of the PPS

**Supplementary Table 3.**
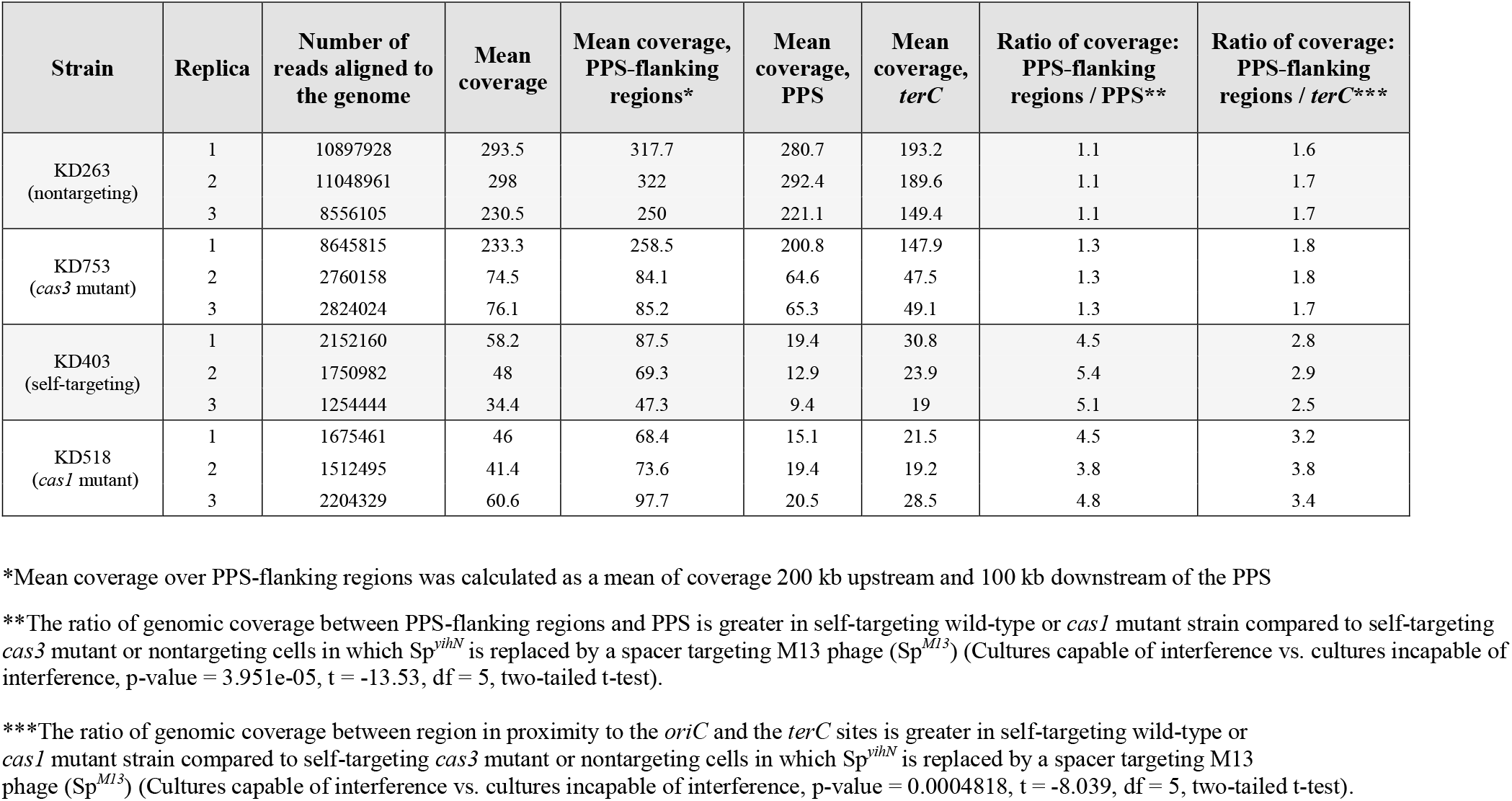
Statistics for sequencing total genomic DNA purified from induced self-targeting strain and control *casl* mutant (Casl H208A), *cas3* mutant (Cas3 H74A) and nontargeting cells. Libraries were prepared only using NEBNext^®^ Ultra™ II DNA Library Prep Kit for Illumina (NEB)

**Supplementary Table 4.** Statistics for sequencing spacers acquired during primed adaptation in self-targeting KD403 strain (number of protospacers 11 each strand upstream or downstream of the PPS)

| Replica | 'Slipped' and 'flipped' AAG-protospacers | Number of newly acquired spacers sequenced | % protospacers from total number of protospacers |  |  |  |
| --- | --- | --- | --- | --- | --- | --- |
|  |  |  | Protospacers in region 100 kb upstream of the PPS |  | Protospacers in region 100 kb downstream of the PPS |  |
|  |  |  | Nontarget strand | Target strand | Nontarget strand | Target strand |
| 1 | Included | 1031147 | 57.3 | 1.2 | 0.8 | 39.8 |
|  | Removed | 1005715 | 57.6 | 0.8 | 0.6 | 40.1 |
| 2 | Included | 1108683 | 57 | 1.2 | 0.9 | 40.3 |
|  | Removed | 1079093 | 57.3 | 0.8 | 0.6 | 40.6 |
| 3 | Included | 1087616 | 56.4 | 1.6 | 1.3 | 38.9 |
|  | Removed | 1058208 | 57 | 1 | 0.9 | 39.3 |

**Supplementary Table 5.**
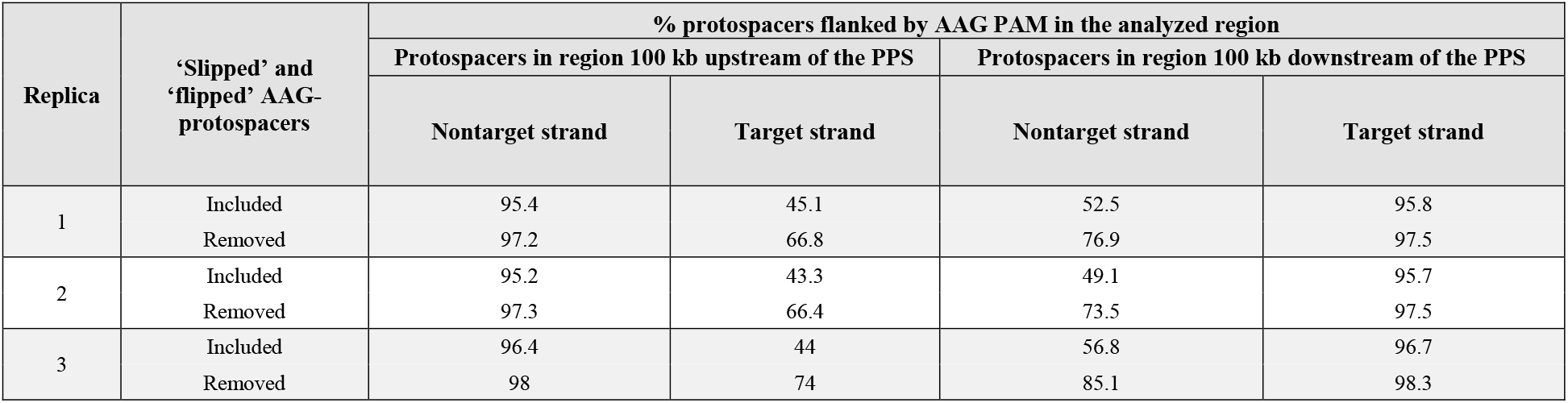
Statistics for sequencing spacers acquired during primed adaptation in self-targeting KD403 strain (% protospacers flanked by AAG PAM on each strand upstream or downstream of the PPS)

| Replica | 'Slipped' and<br>'flipped' AAG-protospacers | % protospacers flanked by AAG PAM in the analyzed region |  |  |  |
| --- | --- | --- | --- | --- | --- |
|  |  | Protospacers in region 100 kb upstream of the PPS |  | Protospacers in region 100 kb downstream of the PPS |  |
|  |  | Nontarget strand | Target strand | Nontarget strand | Target strand |
| 1 | Included | 95.4 | 45.1 | 52.5 | 95.8 |
|  | Removed | 97.2 | 66.8 | 76.9 | 97.5 |
| 2 | Included | 95.2 | 43.3 | 49.1 | 95.7 |
|  | Removed | 97.3 | 66.4 | 73.5 | 97.5 |
| 3 | Included | 96.4 | 44 | 56.8 | 96.7 |
|  | Removed | 98 | 74 | 85.1 | 98.3 |

**Supplementary Table 6.**
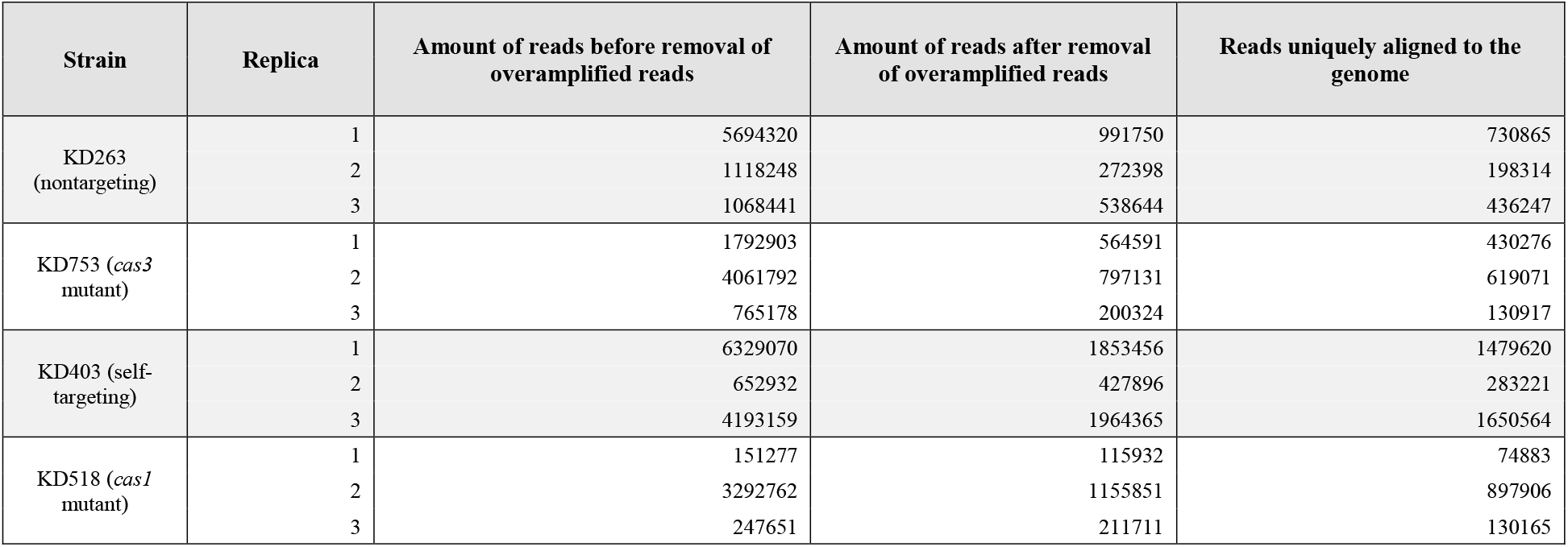
Statistics for sequencing short DNA fragments generated *in vivo* in type I-E system.

**Supplementary Table 7.**
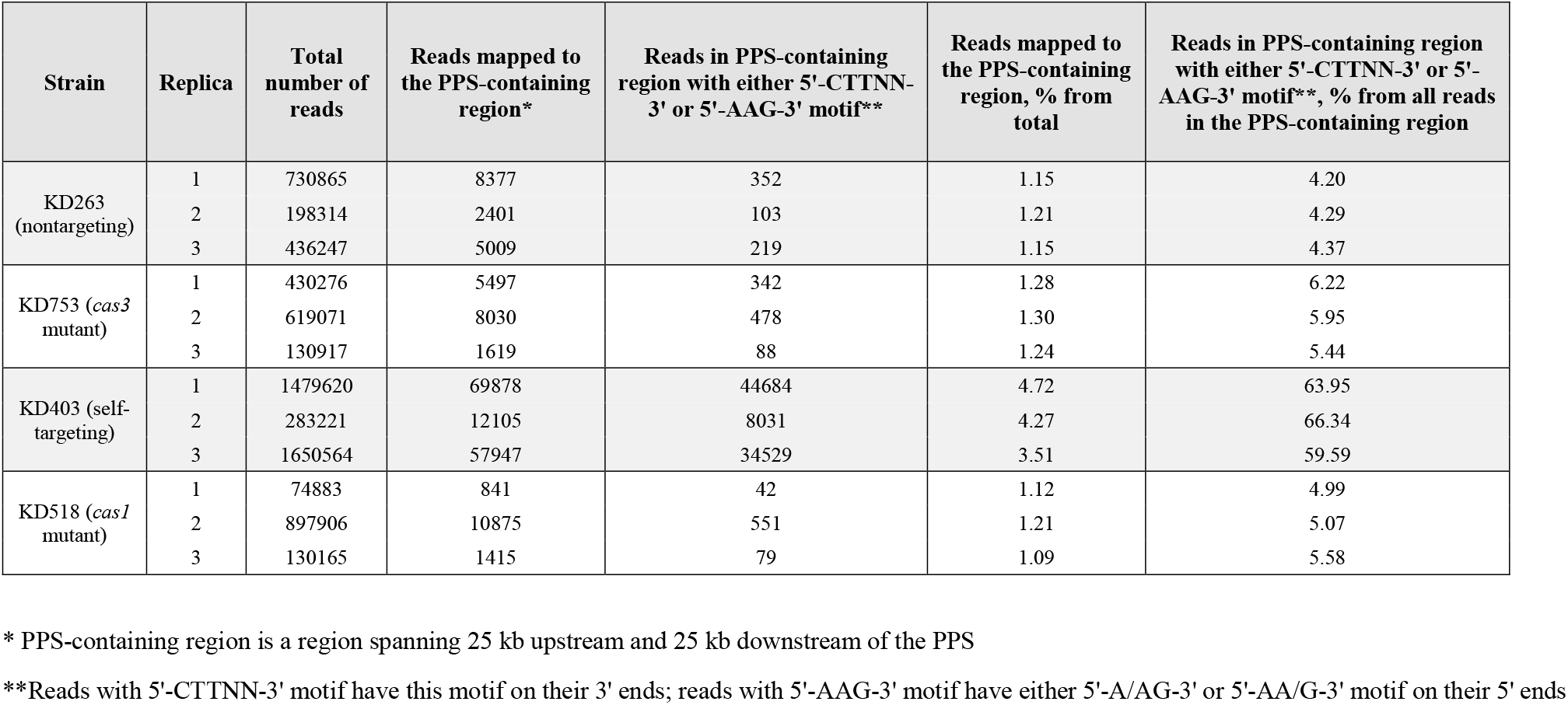
Statistics for sequencing short DNA fragments generated *in vivo* in type I-E system (reads in 50-kb PPS-containing region*)

**Supplementary Table 8.** Statistics for sequencing short DNA fragments generated *in vivo* in type I-E system (reads outside of 50-kb PPS-containing region*)

| Strain | Replica | Total number of reads | Reads mapped outside of the PPS-containing region* | Reads outside of PPS-containing region* with either 5'-CTTNN-3' or 5'-AAG-3' motif** | Reads outside of PPS-containing region with either 5'-CTTNN-3' or 5'-AAG-3' motif**, % from all reads outside of the PPS-containing region* |
| --- | --- | --- | --- | --- | --- |
| KD263 (nontargeting) | 1 | 730865 | 722488 | 27395 | 3.79 |
|  | 2 | 198314 | 195913 | 7084 | 3.62 |
|  | 3 | 436247 | 431238 | 15956 | 3.70 |
| KD753 (*cas3* mutant) | 1 | 430276 | 424779 | 15639 | 3.68 |
|  | 2 | 619071 | 611041 | 21928 | 3.59 |
|  | 3 | 130917 | 129298 | 4696 | 3.63 |
| KD403 (self-targeting) | 1 | 1479620 | 1409742 | 66930 | 4.75 |
|  | 2 | 283221 | 271116 | 12702 | 4.69 |
|  | 3 | 1650564 | 1592617 | 73092 | 4.59 |
| KD518 (*cas1* mutant) | 1 | 74883 | 74042 | 3072 | 4.15 |
|  | 2 | 897906 | 887031 | 32005 | 3.61 |
|  | 3 | 130165 | 128750 | 4803 | 3.73 |

\* PPS-containing region is a region spanning 25 kb upstream and 25 kb downstream of the PPS
\*\*Reads with 5'-CTTNN-3' motif have this motif on their 3' ends; reads with 5'-AAG-3' motif have either 5'-A/AG-3' or 5'-AA/G-3' motif on their 5' ends

**Supplementary Table 9.**
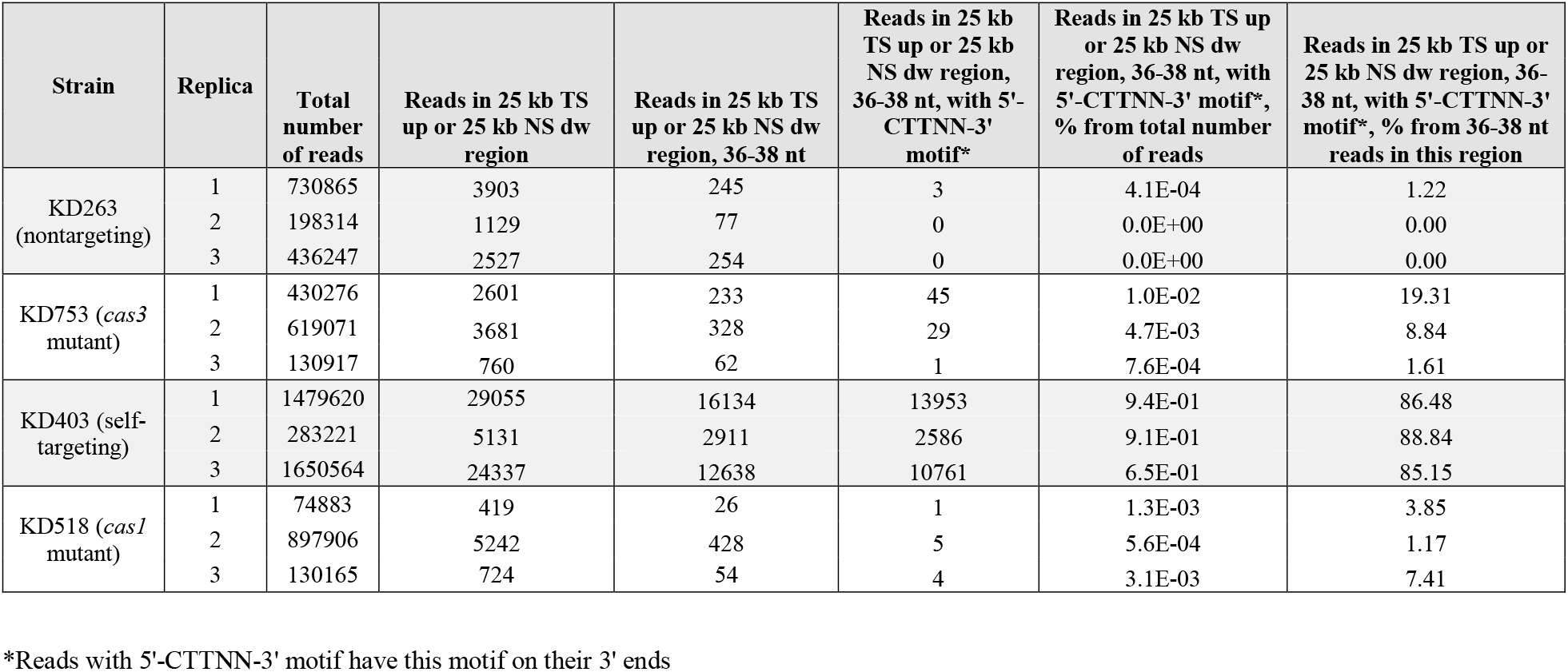
Statistics for sequencing short DNA fragments generated *in vivo* in type I-E system (reads mapped to the target strand in 25kb region upstream of the PPS, “25 kb TS up;” or nontarget strand in 25-kb region downstream of the PPS, “25 kb NS dw”)

**Supplementary Table 10.** Statistics for sequencing short DNA fragments generated *in vivo* in type I-E system (reads mapped to the nontarget strand in 25-kb region upstream of the PPS, “25 kb NS up;” or target strand in 25-kb region downstream of the PPS. “25 kb TS dw”)

| Strain | Replica | Total number of reads | Reads in 25 kb NS up or 25 kb TS dw region | Reads in 25 kb NS up or 25 kb TS dw region, 32-34 nt | Reads in 25 kb NS up or 25 kb TS dw region, 32-34 nt, with 5'-AAG-3' motif* | Reads in 25 kb NS up or 25 kb TS dw region, 32-34 nt, with 5'-AAG-3' motif*, % from total number of reads | Reads in 25 kb NS up or 25 kb TS dw region, 32-34 nt, with 5'-AAG-3' motif*, % from 32-34 nt reads in this region |
| --- | --- | --- | --- | --- | --- | --- | --- |
| KD263 (nontargeting) | 1 | 730865 | 4459 | 246 | 10 | 1.4E-03 | 4.07 |
|  | 2 | 198314 | 1269 | 71 | 2 | 1.0E-03 | 2.82 |
|  | 3 | 436247 | 2477 | 184 | 3 | 6.9E-04 | 1.63 |
| KD753 (*cas3* mutant) | 1 | 430276 | 2867 | 217 | 56 | 1.3E-02 | 25.81 |
|  | 2 | 619071 | 4271 | 320 | 87 | 1.4E-02 | 27.19 |
|  | 3 | 130917 | 839 | 58 | 11 | 8.4E-03 | 18.97 |
| KD403 (self-targeting) | 1 | 1479620 | 40665 | 23868 | 21573 | 1.5E+00 | 90.38 |
|  | 2 | 283221 | 6937 | 4166 | 3846 | 1.4E+00 | 92.32 |
|  | 3 | 1650564 | 33479 | 18777 | 16994 | 1.0E+00 | 90.50 |
| KD518 (*cas1* mutant) | 1 | 74883 | 420 | 54 | 4 | 5.3E-03 | 7.41 |
|  | 2 | 897906 | 5523 | 448 | 96 | 1.1E-02 | 21.43 |
|  | 3 | 130165 | 676 | 53 | 12 | 9.2E-03 | 22.64 |

\*Reads with 5'-AAG-3' motif have either 5'-A/AG-3' or 5'-AA/G-3' motif on their 5' ends

**Supplementary Table 11.** Correlation between 32- to 34-nt 5’-AAG-3’-associated fragments and 36-to 38-nt 5’-CTT-3’ associated fragments in selftargeting KD403 strain.

|  |  |
| --- | --- |
|  | **Fragments with either 5'-A/AG-3' or 5'-AA/G-3' motif on their 5' ends** |
| **Fragments with 5'-CTTNN-3' motif on their 3' ends** | r=0.48 (95% confidence interval: 0.45-0.51); t = 27.667, df = 2538, p-value < 2.2e-16 |

**Supplementary Table 12.** Correlation between number of spacers and corresponding prespacers (DNA fragments conjugated to respective PAM) in self-targeting KD403 strain.

|  | 32- to 34-nt fragments with either 5'-A/AG-3' or 5'-AA/G-3' motif on their 5' ends | 36- to 38-nt fragments with 5'-CTTNN-3' motif on their 3' ends |
| --- | --- | --- |
| Spacers | r=0.57 (95% confidence interval: 0.54-0.59);<br>t = 36.772, df = 2826, p-value < 2.2e-16 | r=0.5 (95% confidence interval: 0.48-0.53);<br>t = 30.976, df = 2826, p-value < 2.2e-16 |

**Supplementary Table 13.**
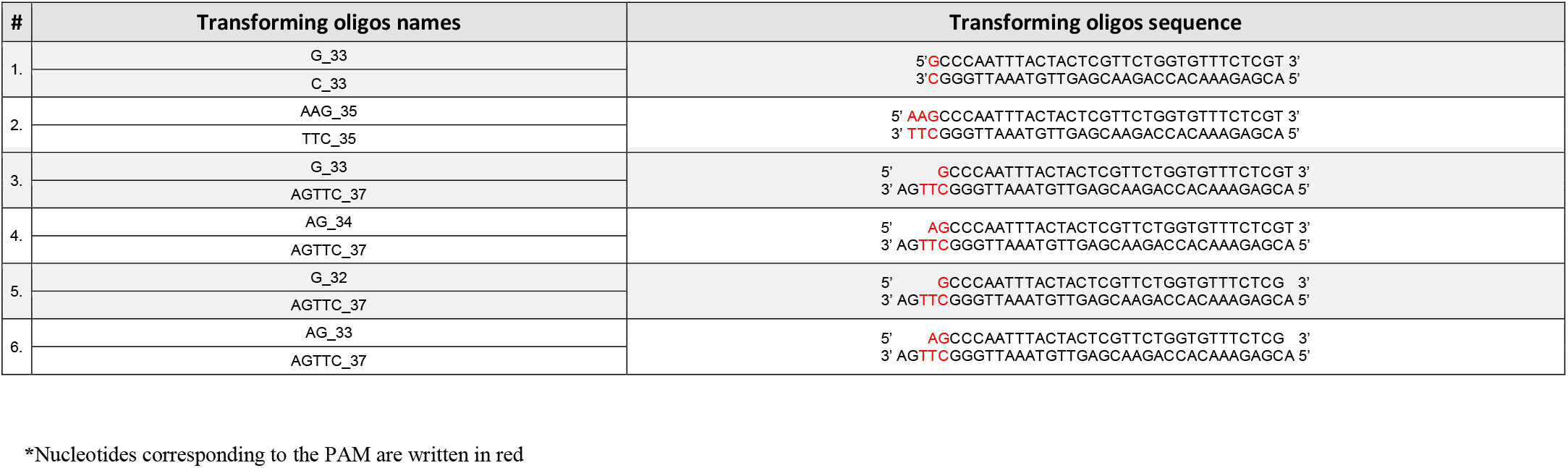
Oligonucleotides used for prespacer efficiency assay.

**Supplementary Table 14.** Oligo transformation assay (overall level of adaptation and source of new spacers)

| Transforming oligo | Replica | Number of CRISPR arrays | CRISPR arrays with an additional spacer | Spacers aligned to genome or pCDF-Cas1/2 | Spacers aligned only to oligo | % of elongated CRISPR arrays | % of CRISPR arrays elongated due to incorporation of oligo-derived spacer | % of CRISPR arrays elongated due to incorporation of a spacer from the genome or pCDF-Cas1/2 |
| --- | --- | --- | --- | --- | --- | --- | --- | --- |
| - | 1 | 527258 | 50183 | 47575 | 49 | 9.52 | 0.01 | 9.02 |
|  | 2 | 595760 | 67484 | 64482 | 42 | 11.33 | 0.01 | 10.82 |
|  | 3 | 763341 | 74111 | 70282 | 66 | 9.71 | 0.01 | 9.21 |
| G_33 + C_33 | 1 | 524532 | 55497 | 48753 | 4058 | 10.58 | 0.77 | 9.29 |
|  | 2 | 384227 | 38315 | 32965 | 3444 | 9.97 | 0.90 | 8.58 |
|  | 3 | 535423 | 49754 | 40736 | 6082 | 9.29 | 1.14 | 7.61 |
| AAG_35 + TTC_35 | 1 | 1226333 | 212389 | 84746 | 116876 | 17.32 | 9.53 | 6.91 |
|  | 2 | 847191 | 170211 | 67824 | 95072 | 20.09 | 11.22 | 8.01 |
|  | 3 | 847361 | 183765 | 54808 | 119865 | 21.69 | 14.15 | 6.47 |
| G_33 + AGTTC_37 | 1 | 953491 | 199266 | 69014 | 119564 | 20.90 | 12.54 | 7.24 |
|  | 2 | 859247 | 186950 | 62169 | 114664 | 21.76 | 13.34 | 7.24 |
|  | 3 | 1056999 | 234441 | 66890 | 152932 | 22.18 | 14.47 | 6.33 |
| AG_34 + AGTTC_37 | 1 | 804648 | 148108 | 48912 | 90757 | 18.41 | 11.28 | 6.08 |
|  | 2 | 768410 | 130580 | 40793 | 82446 | 16.99 | 10.73 | 5.31 |
|  | 3 | 309781 | 55807 | 16765 | 35640 | 18.01 | 11.50 | 5.41 |
| G_32 + AGTTC_37 | 1 | 528490 | 64485 | 38326 | 21154 | 12.20 | 4.00 | 7.25 |
|  | 2 | 496514 | 65306 | 36426 | 23290 | 13.15 | 4.69 | 7.34 |
|  | 3 | 962170 | 141463 | 68424 | 59153 | 14.70 | 6.15 | 7.11 |
| AG_33 + AGTTC_37 | 1 | 626597 | 71983 | 50341 | 16925 | 11.49 | 2.70 | 8.03 |
|  | 2 | 919973 | 88890 | 60539 | 21856 | 9.66 | 2.38 | 6.58 |
|  | 3 | 374155 | 39019 | 25016 | 10813 | 10.43 | 2.89 | 6.69 |

**Supplementary Table 15.**
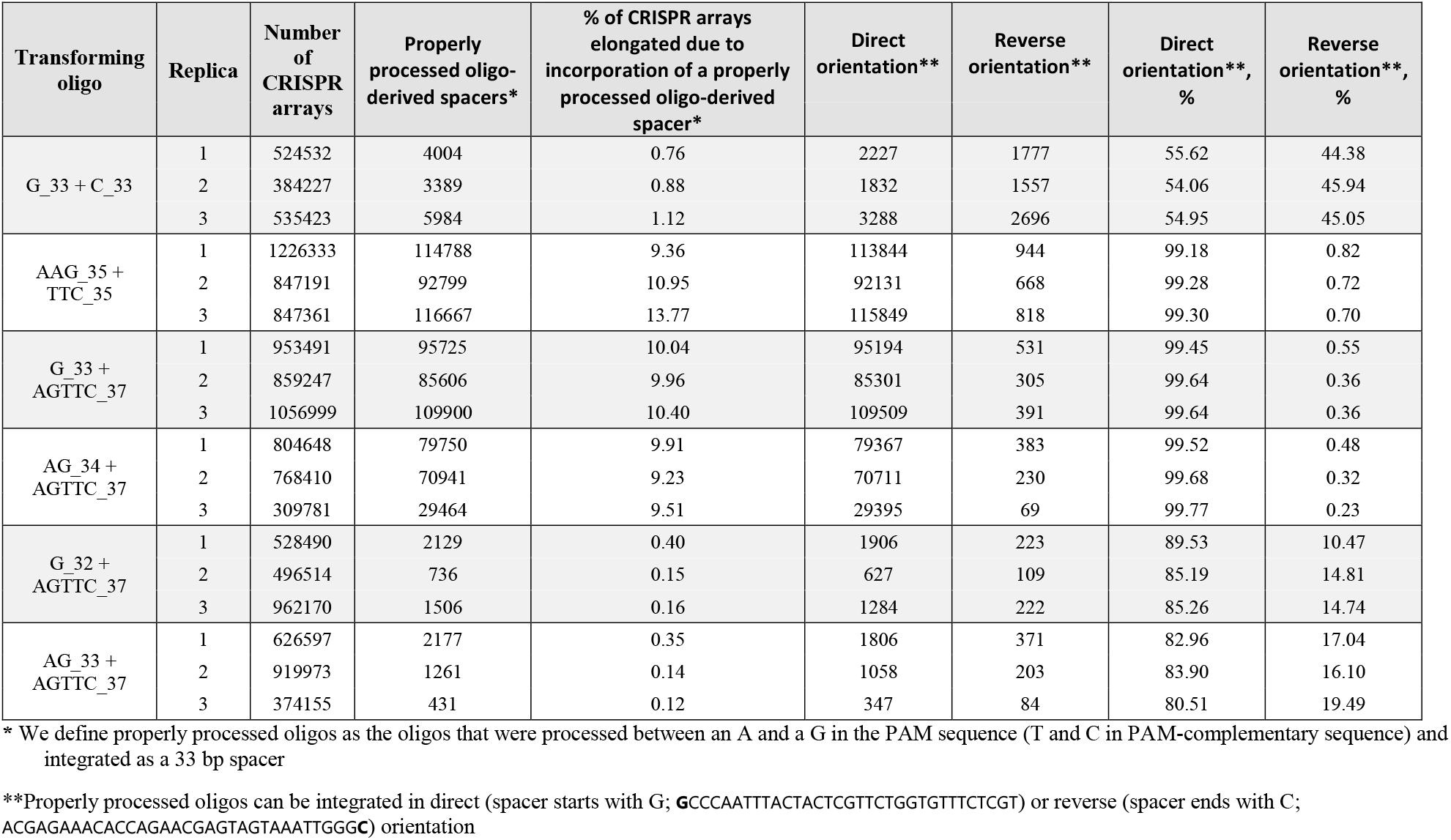
Oligo transformation assay (insertion of properly processed* oligo only)

**Supplementary Table 16.**
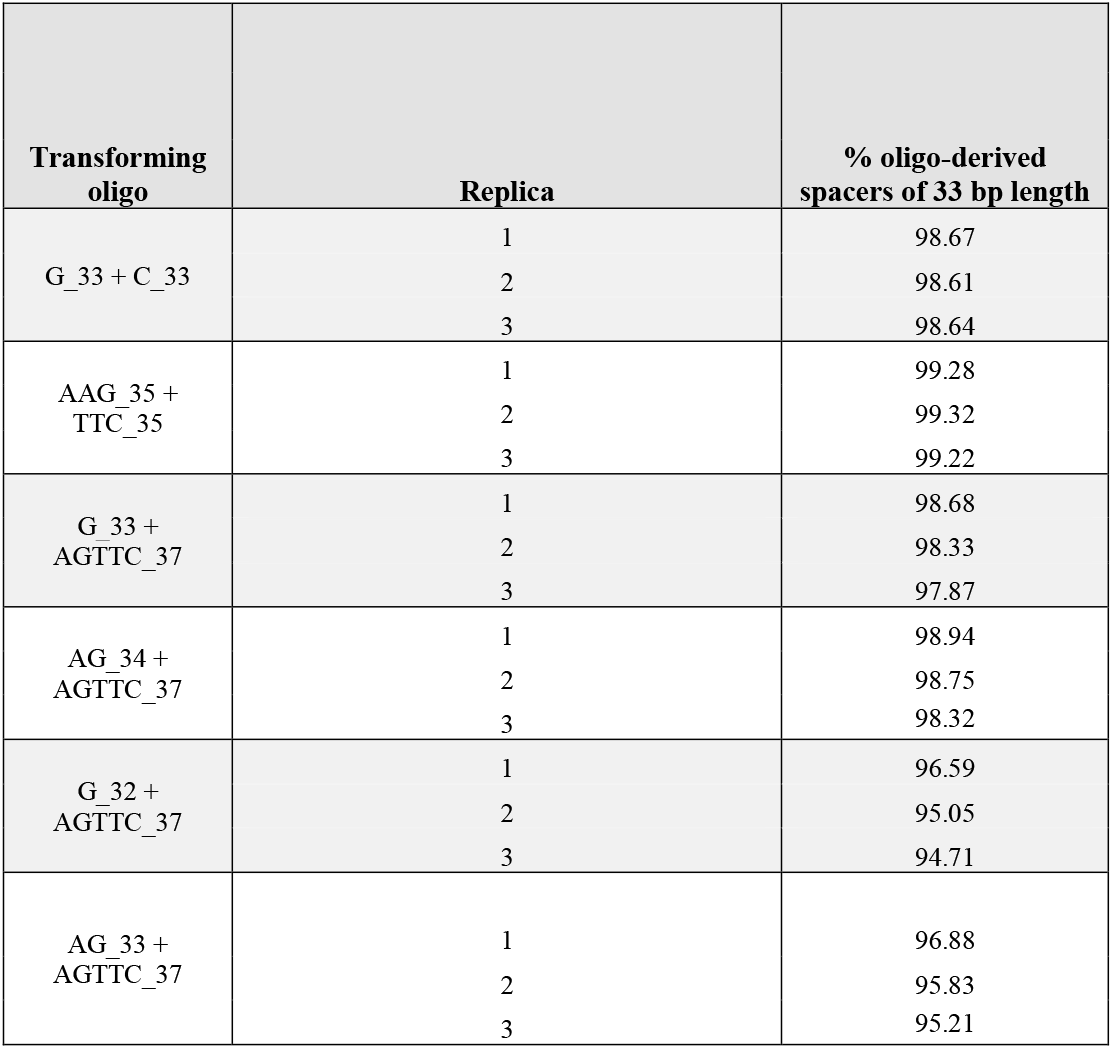
Oligo transformation assay (length of oligo-derived spacers)

**Supplementary Table 17.** Statistics for sequencing short DNA fragments generated *in vivo* in type I-F system.

| Strain | Replica | Amount of reads before removal of overamplified reads | Amount of reads after removal of overamplified reads | Reads uniquely aligned to the genome |
| --- | --- | --- | --- | --- |
| KD675 (*E.coli* strain with type I-F self-targeting system) | 1 | 12437786 | 9739694 | 7034061 |
|  | 2 | 12256224 | 9792344 | 7128753 |

**Supplementary Table 18.** Statistics for sequencing short DNA fragments generated *in vivo* in type I-F system (reads mapped to the target strand in 5kb region upstream of the PPS, “5 kb TS up;” or nontarget strand in 5-kb region downstream of the PPS, “5 kb NS dw”)

| Strain | Replica | Total number of reads | Reads in 5 kb TS up or 5 kb NS dw region | Reads in 5 kb TS up or 5 kb NS dw region, 31-32 nt | Reads in 5 kb TS up or 5 kb NS dw region, 31-32 nt, with 5'-CCA-3' motif* | Reads in 5 kb TS up or 5 kb NS dw region, 31-32 nt, with 5'-CCA-3' motif*, % from 31-32 nt reads in this region |
| --- | --- | --- | --- | --- | --- | --- |
| KD675<br>(*E.coli* strain with type I-F self-targeting system) | 1 | 7034061 | 5740 | 696 | 454 | **65.23** |
|  | 2 | 7128753 | 2665 | 280 | 145 | **51.79** |

\*Reads with 5'-CCA-3' motif have 5'CC/A-3' motif on their 5' ends

**Supplementary Table 19.** Statistics for sequencing short DNA fragments generated *in vivo* in type I-F system (reads mapped to the nontarget strand in 5-kb region upstream of the PPS, “5 kb NS up;” or target strand in 5-kb region downstream of the PPS, “5 kb TS dw”)

| Strain | Replica | Total number of reads | Reads in 5 kb NS up or 5 kb TS dw region | Reads in 5 kb NS up or 5 kb TS dw region, 37-38 nt | Reads in 5 kb NS up or 5 kb TS dw region, 37-38 nt, with 5'-TGGNNN-3' motif* | Reads in 5 kb NS up or 5 kb TS dw region, 37-38 nt, with 5'-TGGNNN-3' motif*, % from 37-38 nt reads in this region |
| --- | --- | --- | --- | --- | --- | --- |
| KD675 (*E.coli* strain with type I-F self-targeting system) | 1 | 7034061 | 5325 | 751 | 260 | **34.62** |
|  | 2 | 7128753 | 2520 | 383 | 123 | **32.11** |

\*Reads with 5'-TGGNNN-3' motif have this motif on their 3' ends

**Supplementary Table 20.** Statistics for sequencing short DNA fragments generated *in vivo* in type I-F system (reads mapped outside of the 10-kb PPS-containing region; 31-32 nt reads)

| Strain | Replica | Total number of reads | Reads outside of the PPS-containing region | Reads outside of the PPS-containing region, 31-32 nt | Reads outside of the PPS-containing region, 31-32 nt, with 5'-CCA-3' motif* | Reads outside of the PPS-containing region, 31-32 nt, with 5'-CCA-3' motif*, % from 31-32 nt reads in this region |
| --- | --- | --- | --- | --- | --- | --- |
| KD675 (*E.coli* strain with type I-F self-targeting system) | 1 | 7034061 | 7022770 | 288270 | 16846 | **5.84** |
|  | 2 | 7128753 | 7123339 | 275324 | 18138 | **6.59** |

\*Reads with 5'-CCA-3' motif have 5'CC/A-3' motif on their 5' ends

**Supplementary Table 21.** Statistics for sequencing short DNA fragments generated *in vivo* in type I-F system (reads mapped outside of the 10-kb PPS-containing region; 37-38 nt reads)

| Strain | Replica | Total number of reads | Reads outside of the PPS-containing region | Reads outside of the PPS-containing region, 37-38 nt | Reads outside of the PPS-containing region, 37-38 nt, with 5'-TGGNNN-3' motif* | Reads outside of the PPS-containing region, 37-38 nt, with 5'-TGGNNN-3' motif*, % from 37-38 nt reads in this region |
| --- | --- | --- | --- | --- | --- | --- |
| KD675 (*E.coli* strain with type I-F self-targeting system) | 1 | 7034061 | 7022770 | 387537 | 9720 | 2.51 |
|  | 2 | 7128753 | 7123339 | 422796 | 9866 | 2.33 |

\*Reads with 5'-TGGNNN-3' motif have this motif on their 3' ends

**Supplementary Table 22.**
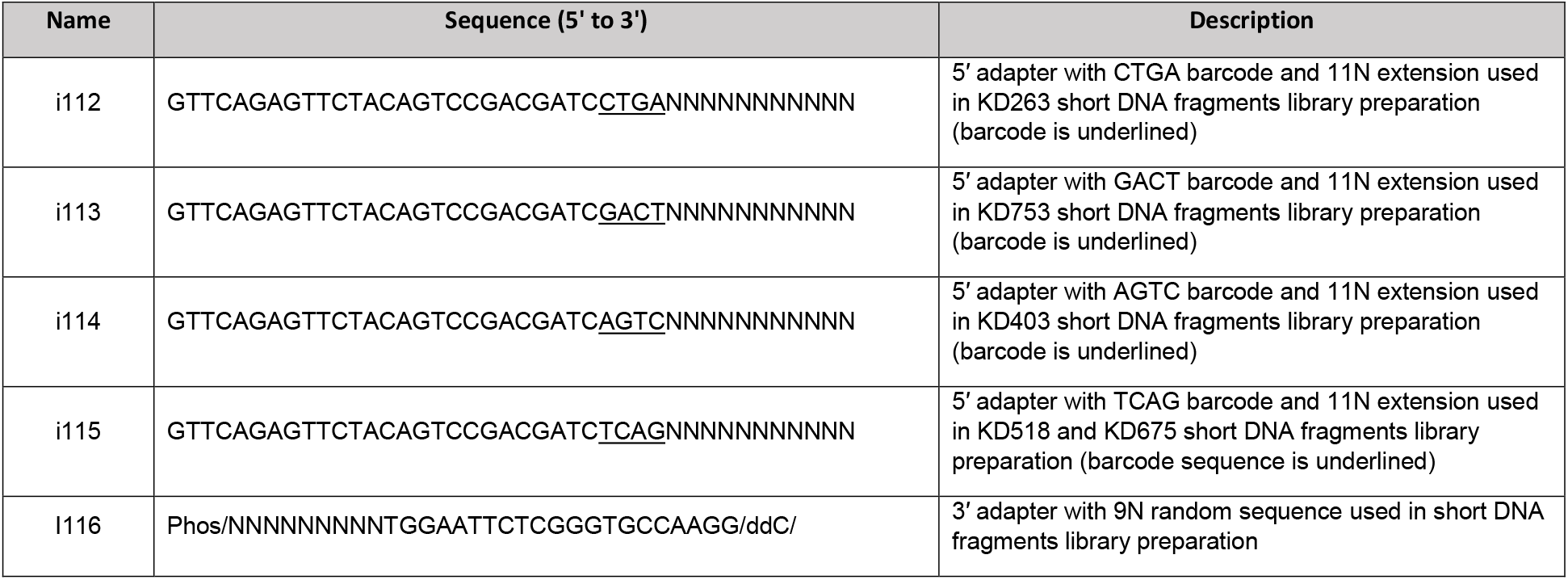
List of adapters used for FragSeq.

**Supplementary Table 23.**
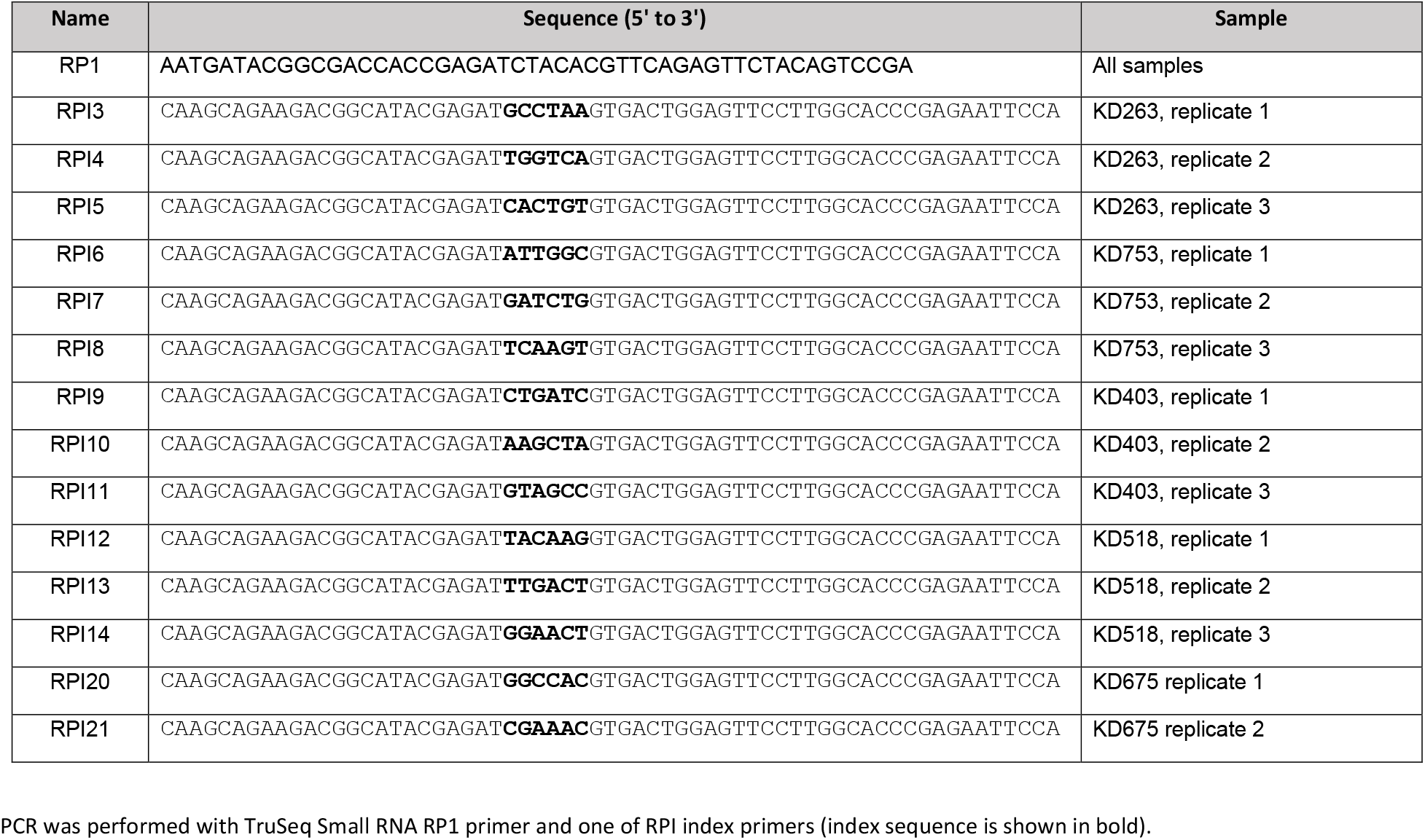
List of Illumina primers used for amplification of FragSeq libraries.

## References

1. Brouns, S. J. et al. Small CRISPR RNAs guide antiviral defense in prokaryotes. Science 321, 960–964, doi: 10.1126/science.1159689 (2008).

2. Datsenko, K. A. et al. Molecular memory of prior infections activates the CRISPR/Cas adaptive bacterial immunity system. Nat Commun 3, 945, doi:10.1038/ncomms1937 (2012).

3. Swarts, D. C., Mosterd, C., van Passel, M. W. & Brouns, S. J. CRISPR interference directs strand specific spacer acquisition. PLoS One 7, e35888, doi:10.1371/journal.pone.0035888 (2012).

4. Yosef, I., Goren, M. G. & Qimron, U. Proteins and DNA elements essential for the CRISPR adaptation process in Escherichia coli. Nucleic Acids Res 40, 5569–5576, doi: 10.1093/nar/gks216 (2012).

5. Mojica, F. J., Diez-Villasenor, C., Garcia-Martinez, J. & Almendros, C. Short motif sequences determine the targets of the prokaryotic CRISPR defence system. Microbiology 155, 733–740, doi: 10.1099/mic.0.023960-0 (2009).

6. Sashital, D. G., Wiedenheft, B. & Doudna, J. A. Mechanism of foreign DNA selection in a bacterial adaptive immune system. Mol Cell 46, 606–615, doi: 10.1016/j.molcel.2012.03.020 (2012).

7. Richter, C. et al. Priming in the Type I-F CRISPR-Cas system triggers strand-independent spacer acquisition, bi-directionally from the primed protospacer. Nucleic Acids Res 42, 8516–8526, doi:10.1093/nar/gku527 (2014).

8. Jore, M. M. et al. Structural basis for CRISPR RNA-guided DNA recognition by *Cascade*. Nat Struct Mol Biol 18, 529–536, doi:10.1038/nsmb.2019 (2011).

9. Wiedenheft, B. et al. Structures of the RNA-guided surveillance complex from a bacterial immune system. Nature 477, 486–489, doi:10.1038/nature10402 (2011).

10. Semenova, E. et al. Interference by clustered regularly interspaced short palindromic repeat (CRISPR) RNA is governed by a seed sequence. Proc Natl Acad Sci USA 108, 10098–10103, doi: 10.1073/pnas.1104144108 (2011).

11. Westra, E. R. et al. CRISPR immunity relies on the consecutive binding and degradation of negatively supercoiled invader DNA by Cascade and Cas3. Mol Cell 46, 595–605, doi: 10.1016/j.molcel.2012.03.018 (2012).

12. Mulepati, S., Orr, A. & Bailey, S. Crystal structure of the largest subunit of a bacterial RNA-guided immune complex and its role in DNA target binding. J Biol Chem 287, 22445–22449, doi: 10.1074/jbc.C112.379503 (2012).

13. Hochstrasser, M. L. et al. CasA mediates Cas3-catalyzed target degradation during CRISPR RNA-guided interference. Proc Natl Acad Sci USA 111, 6618–6623, doi:10.1073/pnas.1405079111 (2014).

14. Hayes, R. P. et al. Structural basis for promiscuous PAM recognition in type I-E Cascade from E. coli. Nature 530, 499–503, doi:10.1038/nature16995 (2016).

15. Mulepati, S. & Bailey, S. In vitro reconstitution of an Escherichia coli RNA-guided immune system reveals unidirectional, ATP-dependent degradation of DNA target. J Biol Chem 288, 22184–22192, doi:10.1074/jbc.M113.472233 (2013).

16. Dillard, K. E. et al. Assembly and Translocation of a CRISPR-Cas Primed Acquisition Complex. Cell 175, 934–946 e915, doi: 10.1016/j.cell.2018.09.039 (2018).

17. Nunez, J. K. et al. Casl-Cas2 complex formation mediates spacer acquisition during CRISPR-Cas adaptive immunity. Nat Struct Mol Biol 21, 528–534, doi: 10.1038/nsmb.2820 (2014).

18. Nunez, J. K., Harrington, L. B., Kranzusch, P. J., Engelman, A. N. & Doudna, J. A. Foreign DNA capture during CRISPR-Cas adaptive immunity. Nature 527, 535–538, doi:10.1038/nature15760 (2015).

19. Nunez, J. K., Lee, A. S., Engelman, A. & Doudna, J. A. Integrase-mediated spacer acquisition during CRISPR-Cas adaptive immunity. Nature 519, 193–198, doi: 10.1038/nature14237 (2015).

20. Savitskaya, E., Semenova, E., Dedkov, V., Metlitskaya, A. & Severinov, K. High-throughput analysis of type I-E CRISPR/Cas spacer acquisition in E. coli. RNA Biol 10, 716–725, doi: 10.4161/rna.24325 (2013).

21. Strotskaya, A. et al. The action of Escherichia coli CRISPR-Cas system on lytic bacteriophages with different lifestyles and development strategies. Nucleic Acids Res 45, 1946–1957, doi:10.1093/nar/gkx042 (2017).

22. Babu, M. et al. A dual function of the CRISPR-Cas system in bacterial antivirus immunity and DNA repair. Mol Microbiol 79, 484–502, doi: 10.1111/j.1365-2958.2010.07465.x (2011).

23. Kunne, T. et al. Cas3-Derived Target DNA Degradation Fragments Fuel Primed CRISPR Adaptation. Mol Cell 63, 852–864, doi:10.1016/j.molcel.2016.07.011 (2016).

24. Shipman, S. L., Nivala, J., Macklis, J. D. & Church, G. M. Molecular recordings by directed CRISPR spacer acquisition. Science 353, aaf1175. doi: 10.1126/science.aaf1175 (2016).

25. Vorontsova, D. et al. Foreign DNA acquisition by the I-F CRISPR-Cas system requires all components of the interference machinery. Nucleic Acids Res 43, 10848–10860, doi: 10.1093/nar/gkv1261 (2015).

26. Redding, S. et al. Surveillance and Processing of Foreign DNA by the Escherichia coli CRISPR-Cas System. Cell 163, 854–865, doi:10.1016/j.cell.2015.10.003 (2015).

27. Nunez, J. K., Bai, L., Harrington, L. B., Hinder, T. L. & Doudna, J. A. CRISPR Immunological Memory Requires a Flost Factor for Specificity. Mol Cell 62, 824–833, doi: 10.1016/j.molcel.2016.04.027 (2016).

28. Wang, J. et al. Structural and Mechanistic Basis of PAM-Dependent Spacer Acquisition in CRISPR-Cas Systems. Cell 163, 840–853, doi:10.1016/j.cell.2015.10.008 (2015).

## References

29. Datsenko, K. A. & Wanner, B. L. One-step inactivation of chromosomal genes in Escherichia coli K-12 using PCR products. Proc Natl Acad Sci U S A 97, 6640–6645, doi: 10.1073/pnas.120163297 (2000).

30. Vischer, N. O. et al. Cell age dependent concentration of Escherichia coli divisome proteins analyzed with ImageJ and ObjectJ. Front Microbiol 6, 586, doi: 10.3389/fmicb.2015.00586 (2015).

31. Morgan, M. et al. ShortRead: a bioconductor package for input, quality assessment and exploration of high-throughput sequence data. Bioinformatics 25, 2607–2608, doi: 10.1093/bioinformatics/btp450 (2009).

32. Okonechnikov, K., Golosova, O., Fursov, M. & team, U. Unipro UGENE: a unified bioinformatics toolkit. Bioinformatics 28, 1166–1167, doi:10.1093/bioinformatics/bts091 (2012).

33. Langmead, B. & Salzberg, S. L. Fast gapped-read alignment with Bowtie 2. Nat Methods 9, 357–359, doi:10.1038/nmeth.1923 (2012).

34. Li, H. et al. The Sequence Alignment/Map format and SAMtools. Bioinformatics 25, 2078–2079, doi:10.1093/bioinformatics/btp352 (2009).

35. Shmakov, S. et al. Pervasive generation of oppositely oriented spacers during CRISPR adaptation. Nucleic Acids Res 42, 5907–5916, doi:10.1093/nar/gku226 (2014).

36. Vvedenskaya, I. O., Goldman, S. R. & Nickels, B. E. Preparation of cDNA libraries for high-throughput RNA sequencing analysis of RNA 5’ ends. Methods Mol Biol 1276, 211–228, doi: 10.1007/978-1-4939-2392-2_12 (2015).

37. Wagih, O. ggseqlogo: a versatile R package for drawing sequence logos. Bioinformatics 33, 3645–3647, doi:10.1093/bioinformatics/btx469 (2017).

